# Pervasive promoter hypermethylation of silenced *TERT* alleles in human cancers

**DOI:** 10.1101/2020.01.29.925552

**Authors:** D Esopi, MK Graham, J Brosnan-Cashman, J Meyers, A Vaghasia, A Gupta, B Kumar, MC Haffner, CM Heaphy, AM De Marzo, AK Meeker, WG Nelson, SJ Wheelan, S Yegnasubramanian

**Author notes:** These authors contributed equally (D Esopi, MK Graham). **Correspondence:** S.J.W.,; S.Y.

## Abstract

In cancers, maintenance of telomeres often occurs through activation of the catalytic subunit of telomerase, encoded by *TERT*. Yet, most cancers show only modest levels of telomerase gene expression, even in the context of activating hotspot promoter mutations (C228T and C250T). The role of epigenetic mechanisms, including DNA methylation, in regulating telomerase gene expression in cancer cells is not fully understood. Here, we have carried out the most comprehensive characterization to date of *TERT* promoter methylation using ultra-deep bisulfite sequencing spanning the CpG island surrounding the core *TERT* promoter in 96 different human cell lines. In general, we observed that immortalized and cancer cell lines were hypermethylated in a region upstream of the recurrent C228T and C250T *TERT* promoter mutations, while non-malignant primary cells were comparatively hypomethylated in this region. However, at the allele-level, we generally observe hypermethylation of promoter sequences in cancer cells is associated with repressed expression, and the remaining unmethylated alleles marked with open chromatin are largely responsible for the observed *TERT* expression in cancer cells. Our findings suggest that hypermethylation of the *TERT* promoter alleles signals transcriptional repression of those alleles, leading to the attenuation of *TERT* activation in cancer cells.

**SIGNIFICANCE:** Hypermethylation of the *TERT* promoter alleles to attenuate *TERT* activation in cancer cells may account for the modest activation of *TERT* expression in most cancers.

## INTRODUCTION

At the terminal ends of human chromosomes are telomere nucleoproteins, consisting of a hexameric DNA repeat, 5’-TTAGGG, coated by a complex of six shelterin proteins (1). Telomeres function as protective caps of chromosomes, safeguarding against genomic instability that can ensue if the exposed ends of chromosomes signaled DNA damage and repair responses. Following each round of cell division, telomeres shorten due to incomplete replication(2,3). Indeed, significant telomere shortening leads to replicative senescence (4), essentially acting as barriers to uncontrolled cell division, while telomere dysfunction is sufficient to induce gross genomic instability (5)(6). Most benign human somatic cells, except for stem cells in rapidly renewing tissues, lack the telomere-specific enzyme telomerase (7), and therefore, are unable to extend and maintain telomeres. However, expressing the catalytic subunit of telomerase, *TERT*, is sufficient for telomerase activity and telomere extension, thus bypassing critical barriers to uncontrolled cell division (8,9).

A defining feature of cancer is limitless replicative potential (10), which for the majority of cancers is facilitated, in part, by the activation of telomerase (11)(7). Highly recurrent mutations (C228T and C250T) in the promoter region of *TERT* can drive *TERT* transcriptional activation and over-expression in cancer (12–14). However, such mutations do not account for the majority of cancers with activated telomerase. In a survey of *TERT* promoter alterations in human cancers, more than 80% of cancers did not harbor *TERT* promoter mutations (15). In these cancers with unaltered *TERT* promoter sequences, it is not always clear how *TERT* expression is activated. Interestingly, regardless of whether *TERT* is activated via promoter mutation or other mechanisms, the degree to which *TERT* is activated appears to be relatively modest, so much so that direct detection of *TERT* mRNA or protein expression using traditional *in situ* methods is uncommon in the majority of cancers (16, 17).

The epigenetic regulation of telomerase expression is not fully characterized, and in particular, the role of DNA methylation of cytosine residues in the control of *TERT* expression remains unresolved. Canonically, promoter DNA methylation is thought to be a repressive chromatin mark, leading to silencing of cis-associated genes. However, there are interesting reports of contexts in which DNA methylation can lead to gene/chromatin activation (18, 19). How DNA methylation may regulate *TERT* expression has been unclear, with multiple conflicting reports. On the one hand, hypermethylation of a small cluster of CpG dinucleotides upstream of the *TERT* transcriptional start site (TSS) was implicated in conferring activation of *TERT* expression in cancer (20,21). However, other studies, including a study examining DNA methylation in cancers with promoter mutations, revealed that *TERT* promoter methylation behaved in the canonical fashion, with promoter methylation associated with silenced alleles, and lack of promoter methylation associated with expressed alleles (22, 23). These conflicting reports may have arisen in part due to the technical challenges in measuring DNA methylation at the TERT promoter regions, which are characterized by very high GC content and CpG density, potentially leading to scant coverage of many relevant CpGs in the regulatory regions using previous methodologies.

Here, we have carried out the most comprehensive characterization of *TERT* promoter methylation to date by developing, extensively validating, and deploying ultra-deep bisulfite sequencing methods to measure DNA methylation at >310 CpGs within and surrounding the core *TERT* promoter in 96 different cell lines, including primary, immortalized, and malignant cells. We find that, generally, hypermethylation of *TERT* promoter sequences in cancer cells is associated with repressed chromatin/expression, and the remaining unmethylated alleles marked with open chromatin marks are largely responsible for the observed *TERT* expression in cancer cells. This association of unmethylated alleles with active expression and methylated alleles with silenced expression is particularly evident in cancers displaying allele specific expression, including cancers harboring recurrent promoter mutations. Overall, these findings suggest that *TERT* promoter hypermethylation in cancer cells may be involved in attenuating the degree of *TERT* activation in cancer cells.

## RESULTS

### Ultra-deep bisulfite sequencing of the TERT promoter CpG island in human cancer and normal cells

The promoter and proximal exons of *TERT* contain a CpG island with particularly high GC content (61.4% G or C, with a ratio of observed to expected CpG of 0.75) (24–27), which has greatly complicated experimental assessment of DNA methylation and other genetic and epigenetic alterations in a comprehensive manner. Whole genome bisulfite sequencing and other genome-scale methodologies often exhibit poor coverage across the entire region, and targeted approaches to overcome such challenges have not comprehensively covered the promoter CpG island around the first exon and upstream regions broadly. Here, we designed and optimized a series of highly overlapping bisulfite sequencing amplicons tiled across both plus and minus strands of the *TERT* promoter CpG island using a microwell, highly-parallel, microfluidics approach. Using this strategy, we were able to overcome these sequencing challenges and characterize the methylation of the *TERT* promoter at nucleotide-level resolution in 96 cell culture models, including 85 cancer lines, 6 immortalized lines, and 5 normal primary cell models **(Table S1)**, as well as a series of reference samples to validate the method.

Notably, the cancer cell lines investigated in this study included both cancer cells with wild type (WT) *TERT* promoters, and cancer cells that harbored highly recurrent *TERT* promoter mutations C228T or C250T. A subset of the cancer cell lines with WT *TERT* promoters employed the telomerase-independent, Alternative Lengthening of Telomeres (ALT) mechanism for telomere maintenance (28). Expression of *TERT*, as assessed by RT-qPCR, confirmed low to no expression in primary and ALT-positive cancer cell models, while immortalized and telomerase-dependent, ALT-negative cancer cell lines had comparably higher *TERT* expression, particularly in the *TERT*-immortalized cell line models **(Figure S1)**.

We interrogated the entire *TERT* promoter region and proximal exons, approximately −1300 base pairs (bp) upstream of the ATG and 2500 bp downstream. Using this bisulfite ultra-deep sequencing approach, we captured over 310 unique CpGs with redundancy by using more than 50 overlapping amplicons on both positive and negative strands **(Figure 1A)**. The methylation measurements of our bisulfite ultra-deep sequencing approach were highly robust, with excellent reproducibility and precision in replicate samples including intra-and inter-batch controls **(Figure S2A)**. Furthermore, our bisulfite ultra-deep sequencing measurements were highly concordant with conventional bisulfite Sanger sequencing measurements **(Figure S2B)**. In our biological controls, methylation measurements were consistent with the known 5-methylcytosine (5mC) genomic content of the well-characterized cell lines, HCT116 DNMT1^KO^, HCT116 DNMT3b^KO^, and HCT116 DKO, which is deficient in both *DNMT1* and *DNMT3b*. HCT116 DKO cells, known to harbor approximately 5% of the 5mC content of HCT116 WT cells (29), showed significant hypomethylation compared to WT parental HCT116 cells. Additionally, HCT116 cell line models deficient in either *DNMT1* or *DNMT3b*, each known to harbor 60% and 80% reduced 5mC content compared to WT parental cells (29), showed concordant levels of demethylation around the *TERT* promoter in our bisulfite ultra-deep sequencing data **(Figure S2C)**. Taken together, these analyses established that our approach provided robust and accurate methylation measurements across the *TERT* promoter.

**Figure 1.**
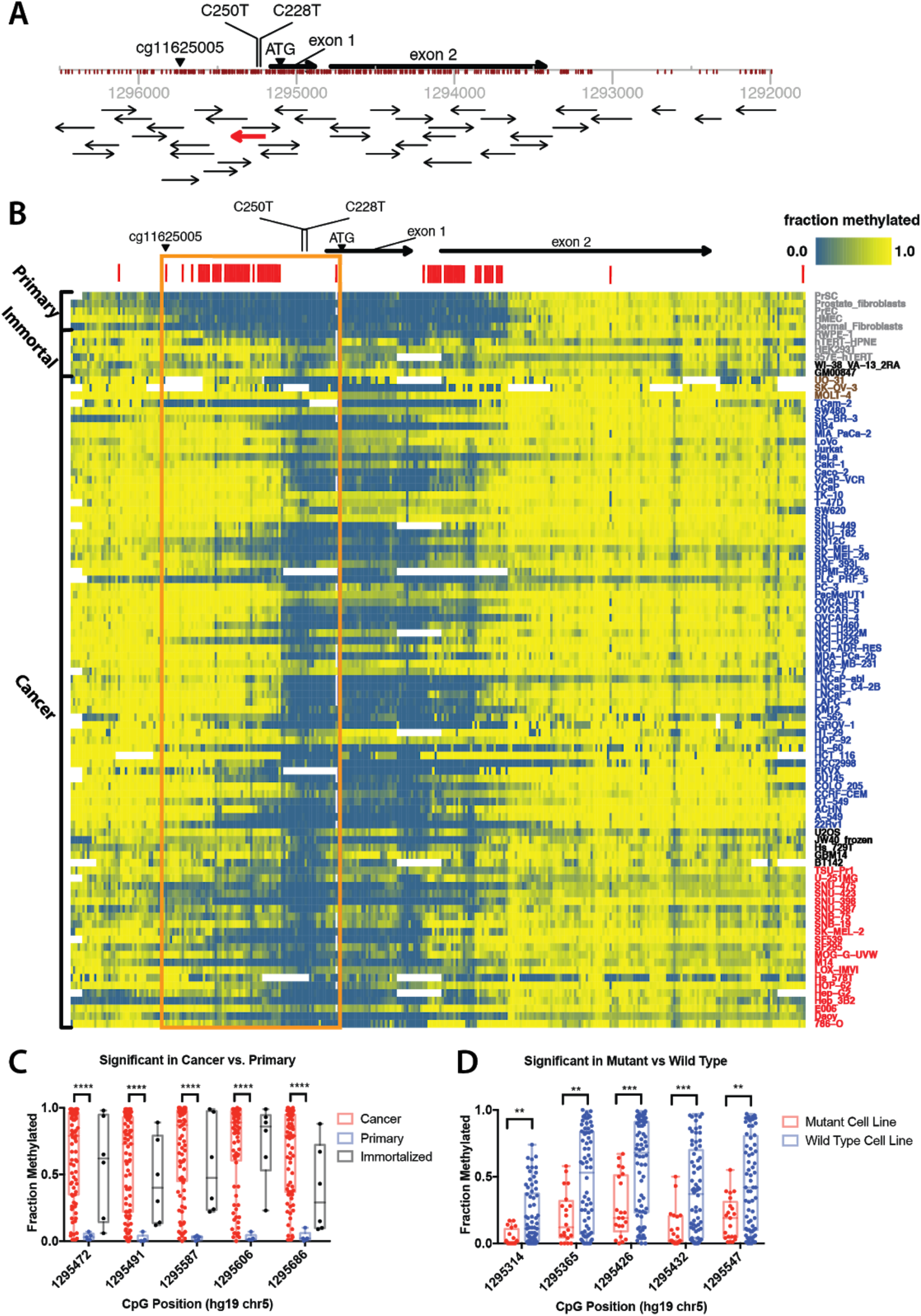
Bisulfite deep sequencing of the *TERT* promoter. Comprehensive characterization of *TERT* promoter methylation using ultra-deep bisulfite sequencing of >310 CpGs within and surrounding the core *TERT* promoter in 96 different cell lines – 85 cancer cell models, 6 immortalized cell models, 5 normal cell models in primary culture. (A) Schematic of the *TERT* promoter locus with position and orientation of all *TERT* Access Array amplicons, the positions of all interrogated CpGs, and notable landmark positions (ATG, C228T and C250T mutations, and cg11625005 Infinium probe location). Amplicon highlighted in red overlaps the highly recurrent C228T and C250T mutations. (B) Characterization of the methylation pattern of the *TERT* promoter at nucleotide-resolution shows that a region of CpGs in the core promoter and upstream of transcriptional start site (TSS) are generally hypermethylated in cancers compared to normal samples (orange box). Red bars above the heatmap denote the top 20% most variable CpGs across all samples. Cell line names are colored based on status (grey = primary and immortalized cell lines, black = ALT-positive cell lines, blue = cancer cell lines with WT *TERT* promoter, red = cancer cell lines with mutant TERT promoter, brown = cancer cell lines with unknown *TERT* promoter mutation status). (C) Top 5 CpGs most differentially methylated between cancer and primary cells. Plot shows fraction methylation of CpGs in cancer, primary, and immortalized cell lines. (D) Top 5 CpGs most differentially methylated between cancers with WT *TERT* promoters and mutated TERT promoters. Mann-Whitney test was used to assess statistically significant difference. Asterisks denote level of significance (* p ≤ 0.5, ** p ≤ 0.01, *** p ≤ 0.001, **** p ≤ 0.0001).

In our expanded analysis of 96 different samples, we observed broadly that multiple large regions far upstream (chr5:1295338-1295731, hg19) and downstream (chr5:1294565-1294844, hg19) of the highly recurrent C228T and C250T mutations were nearly ubiquitously methylated **(Figure 1B)**. The top 20% most variably methylated CpGs across all samples occurred in two major regions, one upstream of the location of the recurrent promoter mutations, and another downstream of exon 1, with high correlation of methylation within each of these regions **(Figure S3A)**. Clustering samples by the top 20% most variable CpGs revealed that a cluster containing all of the non-malignant primary cells generally harbored less methylation than cancer cells **(Figure S3B; Figure 1B)**. Indeed, identifying differentially methylated CpGs between cancer and normal samples indicated that, generally, the differentially methylated CpGs were more methylated in cancer compared to normal (see rightward skew in **Figure S4A)**, with a significant fraction of these, including the top five, occuring in a region upstream of the hotspot mutations **(Figure S3B, Figure 1C)**.

We further validated these differential methylation patterns between cancer and non-malignant samples using previously published methylation microarray data. We compared our methylation assessment of a CpG residue that was also interrogated by the Infinium Methylation microarray platform (Infinium probe cg11625005, chr5:1295737, hg19) upstream of the TSS, which has been implicated as a marker of malignancy in cancer (20,30,31). In our panel of cell line models, we found that cancer cells were generally more methylated than normal cells at this CpG. Similarly, analysis of the publicly available TCGA data (32) confirmed that cancers were typically more methylated than normal at this CpG **(Figures S4B-C)**.

### Allele-Specific hypermethylation of wild-type alleles in C228T/C250T promoter mutant cancers

We next identified CpGs that were differentially methylated between WT and *TERT* promoter mutant cancers. Interestingly, in general, the differentially methylated CpGs, including the top 5 CpGs, showed hypomethylation in the mutant compared to WT cancers, with many of these positions occurring just upstream of the position of the recurrent promoter mutations **(Figure S5A; Figure 1D)**.

Given the proximity of these differentially methylated CpGs in WT and mutant cancers to the position of the promoter mutations, we next assessed whether the mutant cancers exhibited allele-specific methylation of mutant or WT alleles. Although C -> T mutations can be difficult to assess in bisulfite sequencing data, we took advantage of the fact that we designed overlapping amplicons directed to the plus and minus strand separately and identified an amplicon (red arrow, **Figure 1A**) that interrogated the G -> A mutation on the complementary strand, which would not be affected by bisulfite conversion. This amplicon allowed us to assess the mutation status of each sequenced allele, and also allowed phasing of the surrounding CpG methylation patterns with WT or mutant alleles (**Figure S5B**). Notably, this allowed, for the first time, phasing of CpG methylation to mutational status at single molecule resolution for each allele. Sanger sequencing of genomic DNA from a subset of the cell lines assessed in our study confirmed that our approach could accurately identify TERT promoter mutations from the bisulfite sequencing data **(Figure S5C)**. Our deep sequencing data from this amplicon revealed that nearly all promoter mutant cancers showed allele specific methylation of the WT allele and significantly reduced methylation of the mutant allele in a region upstream of the mutations **(Figure 2A,B)**. Six neighboring CpGs were found to be most differentially methylated in the amplicon containing highly recurrent *TERT* promoter mutations **(Figure 2A)**. Interestingly, the distribution of methylation of the WT alleles in mutant cell lines was similar to that of WT cell lines, with mutant alleles in the mutant cell lines being significantly hypomethylated **(Figure 2B, Figure S4A, Figure S5A)**.

**Figure 2.**
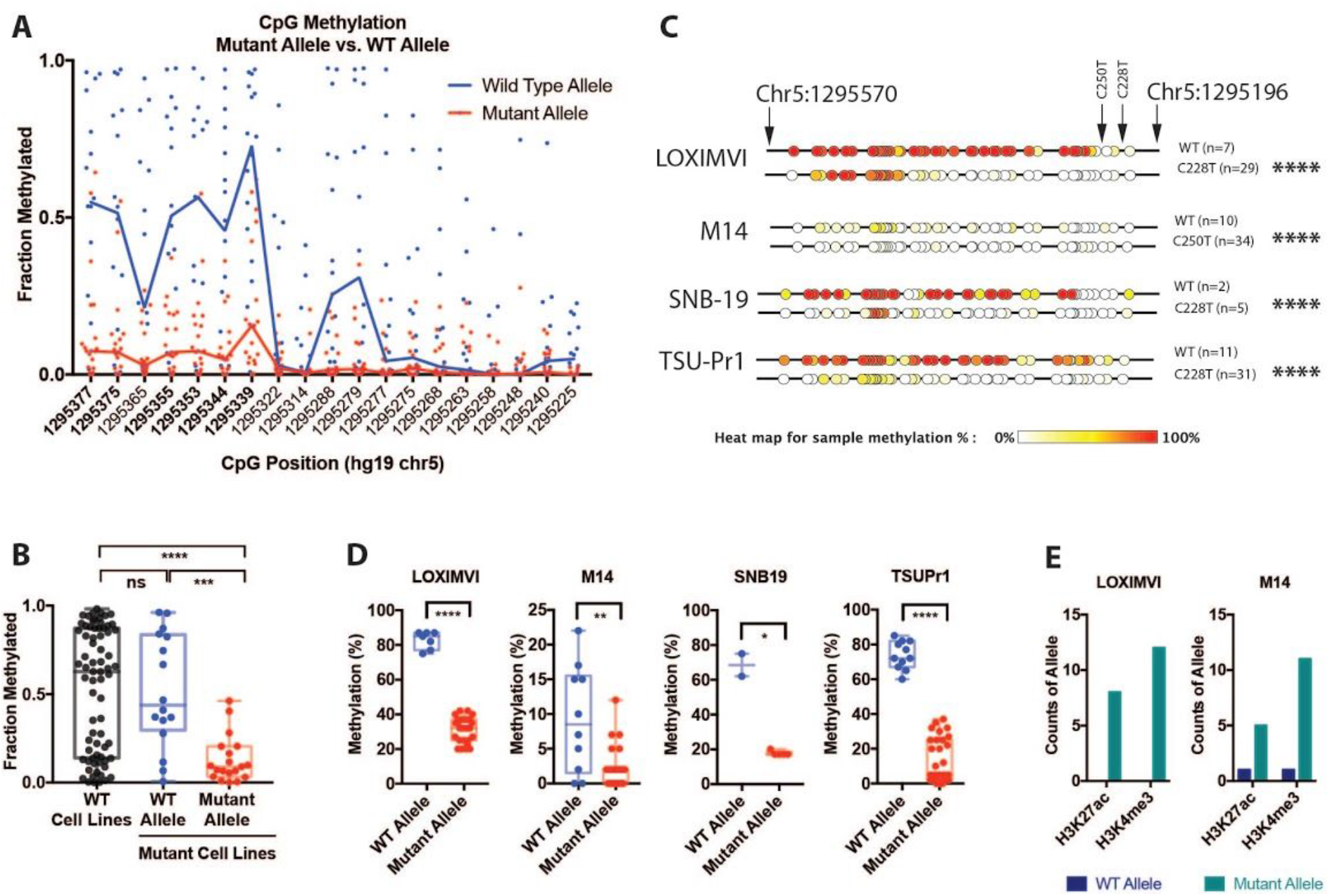
Allele-specific methylation in cancer cells with recurrent mutations in *TERT* promoter region. Sequenced alleles were phased to assess CpG methylation patterns upstream of the recurrent promoter mutations C228T and C250T. (A) A scatter distribution plot of mutant and WT alleles shows allele-specific hypermethylation of the WT allele and reduced methylation of the mutant allele in a region upstream of the mutations, particularly at 6 CpGs (positions bolded in plot). (B) Methylation of the 6 differentially methylated CpGs were compared between WT and mutant alleles in cancer cells harboring *TERT* promoter mutations. Generally, mutant alleles were hypomethylated, while the WT alleles of mutant cell lines and alleles from WT cell lines were hypermethylated. (C) CpG methylation maps and (D) distribution plots from conventional bisulfite Sanger sequencing of cancer cell lines with *TERT* promoter mutations, LoX-IMVI, M14, SNB19, and TSU-PR1 validate bisulfite deep sequencing results, showing WT alleles are hypermethylated and mutant alleles are hypomethylated. (E) ChIP of LoX-IMVI and M14 cells show that mutant alleles are enriched with the open chromatin marks, H3K4me^3^ and H3K27ac. Mann-Whitney test was used to assess a statistically significant difference in plots, while Chi-square test was used in a 2×2 contingency table of *TERT* promoter status of allele (WT vs mutant) and GpG methylation (methylated vs unmethylated). Asterisks denote level of significance (* p ≤ 0.5, ** p ≤ 0.01, *** p ≤ 0.001, **** p ≤ 0.0001).

To investigate this allele-specific methylation in greater detail, we selected representative cell lines with promoter mutations, and performed bisulfite Sanger sequencing of a larger region that could interrogate an increased number of CpGs upstream of the promoter mutations. These analyses confirmed allele-specific hypomethylation of mutant alleles compared to WT alleles in cancer cell line models with C228T or C250T *TERT* promoter mutations, namely, LOX-IMVI, M14, SNB19, and TSU-PR1 cells **(Figure 2C)**. Furthermore, the allele-specific methylation patterns of the six differentially methylated CpGs identified in the deep sequencing data were also found to be significantly differentially methylated between WT and mutant alleles in the bisulfite Sanger sequencing in these representative promoter mutant cell lines **(Figure 2D)**.

We assessed whether the reduced methylation seen in the mutant alleles compared to WT alleles translated to establishment of chromatin marks associated with active chromatin. In LOX-IMVI and M14 cells, we performed chromatin immunoprecipitation (ChIP) for epigenetic marks of active chromatin, H3K4me^3^ and H3K27ac, followed by qPCR and Sanger sequencing of the products. These analyses revealed that mutant alleles were enriched for DNA hypomethylation, increased H3K4me^3^, and increased H3K27ac, while WT alleles were hypermethylated **(Figure 2D,E)**. In the context of previous studies showing strong allele-specific expression of mutant alleles and lack of expression of WT alleles in mutant cancers(33,34), our results suggest that DNA hypermethylation is associated with transcriptionally silent WT alleles, while DNA hypomethylation is associated with active chromatin marks and transcription from mutant alleles.

### Rare cell lines with predominantly mutant TERT promoters still maintain a balance of methylated repressed and unmethylated active alleles

It has been suggested that mutant *TERT* promoter alleles are undermethylated because they can prevent the polycomb complex and epigenetic machinery from promoting methylation of the adjacent sequence (22). If this were the case, we would expect that the few cancers that have bi-allelic or hemizygous promoter mutations should be profoundly undermethylated in all alleles. To examine this possibility, we used our deep sequencing data to identify four cell lines with >97% mutant allele fraction for the C228T or C250T mutations, indicating bi-allelic or hemizygous mutation: U251, HOP62, SK-MEL-2, and SF539. Sanger sequencing of genomic DNA confirmed that indeed U251 and HOP62 harbor predominantly mutant *TERT* promoter alleles **(Figure S7)** validating the deep sequencing data in accurately identifying cell lines with bi-allelic/hemizygous promoter mutations.

Of these four cell lines, SK-MEL2 and SF539 had >80% of alleles with very low methylation as would be expected, a finding confirmed by bisulfite Sanger sequencing **(Figure S6)**. However, the other 2 cell lines, HOP62 and U251, harbored >55% of the mutated alleles showing increased methylation upstream of the mutation sites, with the remaining mutant alleles showing low methylation levels similar to that seen in the mutant alleles of cancers with mono-allelic mutation (**Figure 3**). Subsequent bisulfite Sanger sequencing of SK-MEL-2 and HOP62 confirmed the contrasting methylation patterns identified in the deep sequencing data (**Figure 3B, Figure S6**).

**Figure 3.**
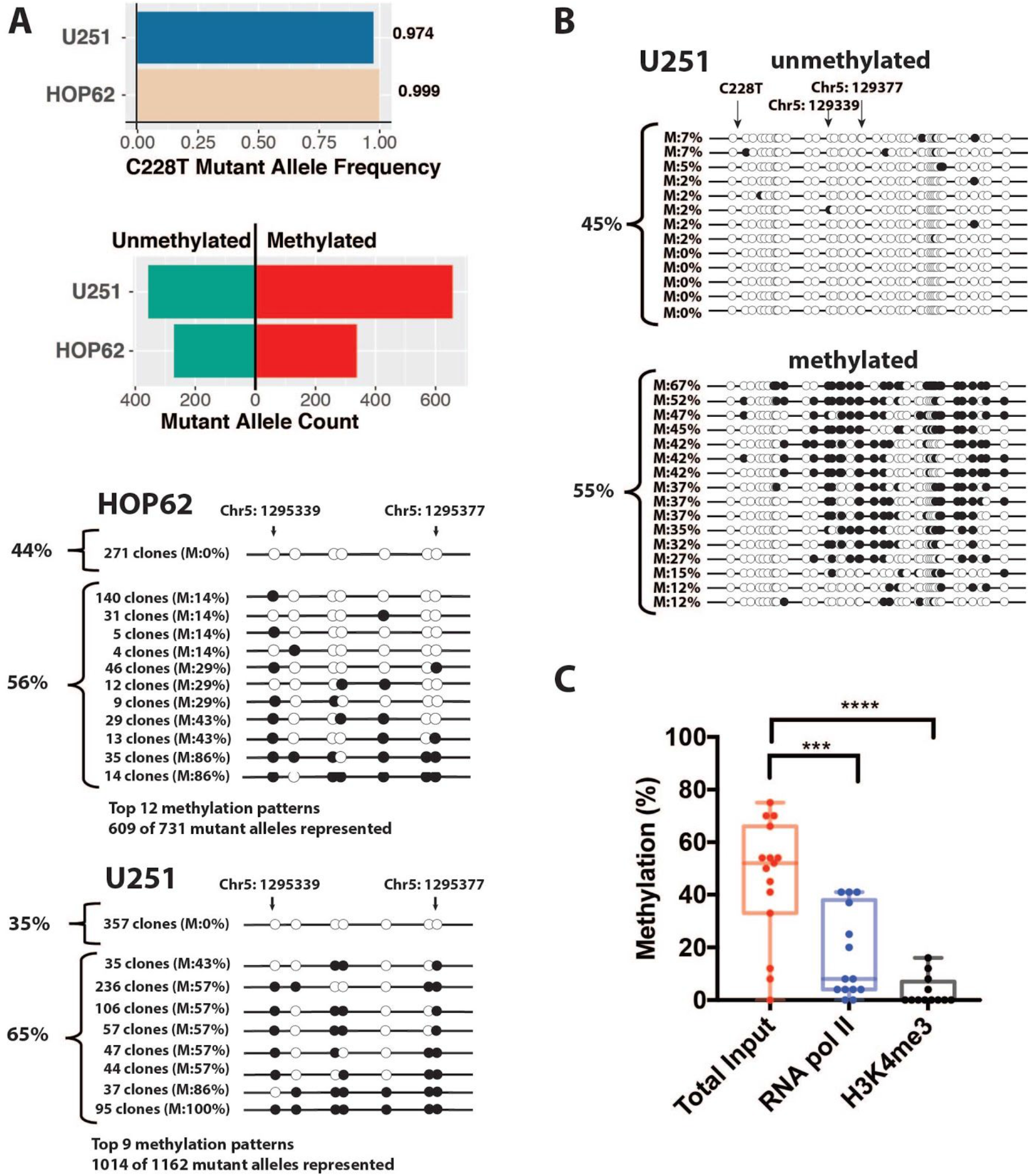
Rare cancer cells with high mutant allele fractions maintained methylated repressed and unmethylated active alleles. Cell lines with high mutant allele fractions had a mix of methylated repressed and unmethylated active mutant alleles. (A) Mutant cancer cell lines HOP62 and U251 had predominantly mutant alleles, but a balance of methylated and unmethylated alleles, as assessed by bisulfite deep sequencing. Black circles indicate methylated CpGs and open circles indicate unmethylated CpGs. (B) Conventional bisulfite Sanger sequencing of U251 validated bisulfite deep sequencing results. (C) ChIP of U251 showed that unmethylated alleles were enriched for RNA pol II occupancy, and the open chromatin mark H3K4me^3^. Mann-Whitney test was used to assess a statistically significant difference. Asterisks denote level of significance (* p ≤ 0.5, ** p ≤ 0.01, *** p ≤ 0.001, **** p ≤ 0.0001).

HOP62 and U251 thus represent interesting exceptions to the general trend that mutant alleles are undermethylated, suggesting that the presence of a mutation does not automatically lead to protection from DNA methylation. We hypothesized that, like other cancers with heterozygous mutations, the unmethylated and mutated alleles should be enriched for chromatin activation marks and be more selectively bound by the transcriptional machinery compared to the methylated and mutated alleles. To test this hypothesis, we carried out ChIP followed by qPCR and bisulfite sequencing in U251 cells. After confirming that the histone activation mark H3K4me3 and RNA Polymerase 2 (Pol2) were both highly enriched at the U251 *TERT* promoter using ChIP-qPCR (**Figure S8**), we found that both of these chromatin activation factors were highly enriched for the unmethylated alleles using ChIP bisulfite sequencing **(Figure 3C)**. Taken together, these data suggest that some cancers with bi-allelic or hemizygous *TERT* promoter mutations still maintain a high fraction of methylated alleles, with only the remaining unmethylated alleles being associated with active chromatin and RNA Pol 2 binding.

### Cells without promoter mutation show hypermethylation of inactive alleles and hypomethylation of active alleles

In cancer cell lines with C228T/C250T promoter mutations and monoallelic expression, the mutant allele is expressed while the WT allele is repressed (33,34). However, cancer cell lines can also have monoallelic expression of *TERT* even in the absence of the C228T/C250T promoter mutations (34). We examined, in cancers with monoallelic expression and WT *TERT* promoters, whether the non-expressed allele was selectively methylated. Among the cell lines analyzed in our sample set, RPMI-8226 was identified to have monoallelic expression. Sanger sequencing of genomic DNA confirmed that RPMI-8226 was indeed heterozygous for both A and G alleles of the rs2736098 SNP located in exon 2 of *TERT* **(Figure 4A)**, while Sanger sequencing of the cDNA derived from the TERT mRNA confirmed monoallelic expression of only the A allele **(Figure 4B)**. We performed long-range bisulfite amplicon Sanger sequencing (~1.5 kb), which encompassed the core *TERT* promoter, the region upstream of the highly recurrent promoter mutations, and the heterozygous rs2736098 SNP in order to phase the promoter DNA methylation pattern with the active vs. inactive allele **(Figure S9)**. We observed that the expressed A allele phased with unmethylated sequences, while the repressed G allele phased with methylated sequences **(Figure 4C)**. The average methylation of the six most differentially methylated CpGs between mutant and WT alleles upstream of the highly recurrent *TERT* promoter mutations, originally identified in our deep sequencing data, showed significant difference between the A and G alleles, with the expressed A allele being hypomethylated and the repressed G allele being hypermethylated **(Figure 4D)**. This interesting example of a cancer with mono-allelic *TERT* expression provides further evidence that *TERT* promoter DNA methylation is associated with inactive alleles.

**Figure 4.**
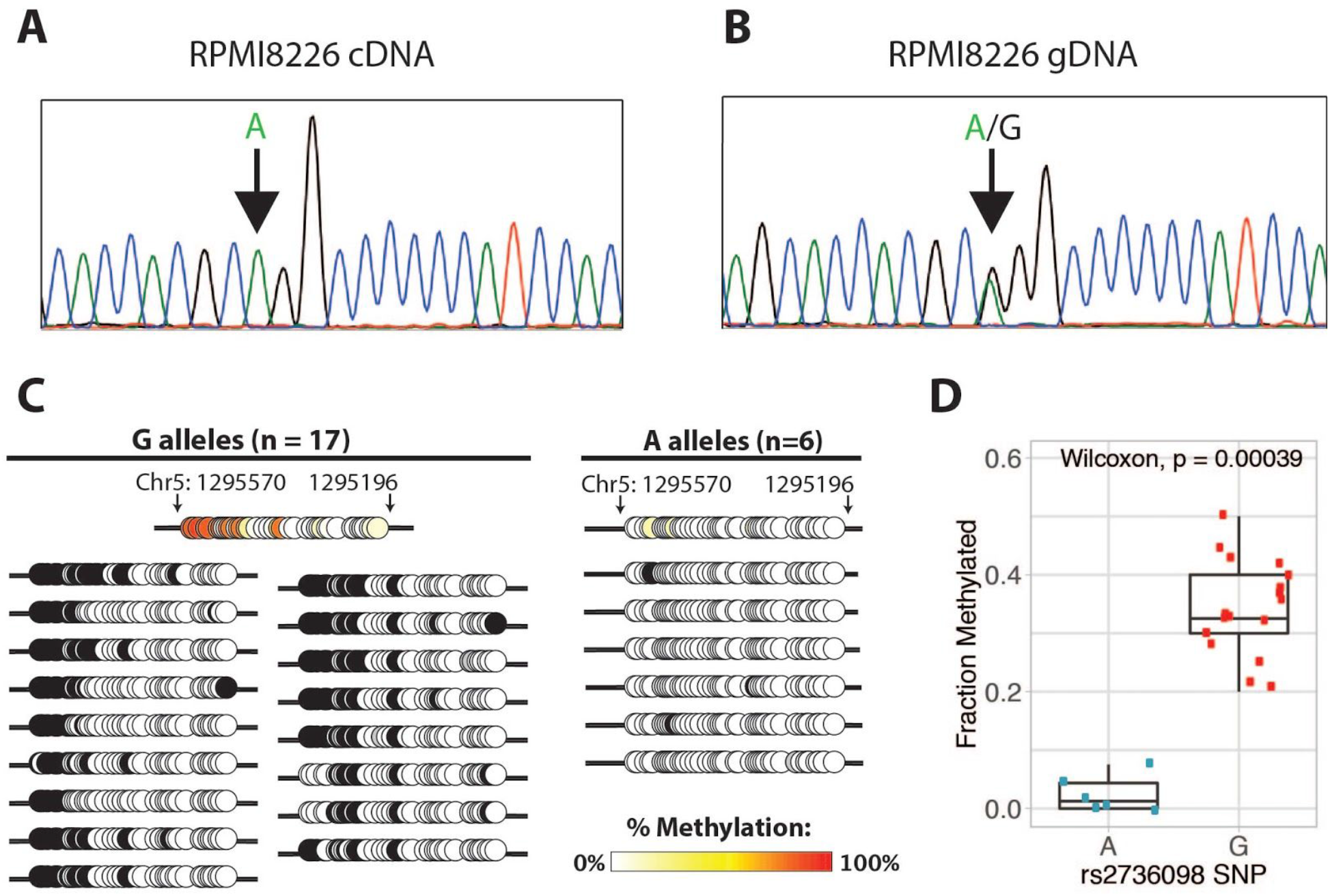
Hypermethylation of the repressed allele and hypomethylation of the active allele in WT promoter. Monoallelic expression of *TERT* from the cell line RPMI8226 demonstrated selective expression of the hypomethylated allele. Sanger sequencing of (A) cDNA and (B) gDNA in the rs2736098 locus revealed that RPMI8226 possessed both the A and G alleles, but expressed only the A allele. (C) CpG methylation maps from long-range bisulfite sequencing, showing only the region overlapping the highly recurrent C228T and C250T mutations in the TERT promoter region hg19 chr5: 1295196 - 1295570. Expressed allele A is unmethylated while the unexpressed G allele is methylated. Black circles indicate methylated CpGs and open circles indicate unmethylated CpGs. (D) The box plot includes the median, 25% quantile, and 75% quantile of fraction methylated for A and G alleles. The whiskers show the 95% confidence interval for the median.

### Methylation of TERT promoter sequences results in strong repression of heterologous reporter constructs

Finally, we assessed whether methylation of the *TERT* promoter, whether WT or mutant, would silence expression in a luciferase construct in multiple cell models, including LNCaP (TERT WT cancer cell line), SNB-19 (TERT mutant cancer cell line), U2OS (telomerase-negative, ALT-positive cancer cell line), and HEK293T (immortalized cell line). Multiple deletion constructs of the *TERT* promoter were treated with the methyltransferase enzyme *M. SssI* or mock-treated to produce one set of fully methylated and one set of fully unmethylated deletion constructs, respectively **(Figure 5A)**. In these studies, a reporter construct driven by the glutathione S-transferase Pi 1 (*GSTP1*) promoter were also included. The *GSTP1* promoter is overlapped by a well characterized CpG island, and methylation of this promoter is known to silence gene expression (35). HEK293T cells transfected with various deletion constructs of the *TERT* promoter driving luciferase expression showed activity similar to the control reporter construct driven by the *GSTP1* promoter. The only exceptions were the deletion constructs that lacked elements of the core promoter, which showed reduced activity **(Figure 5B)**. As expected, methylation of the *GSTP1* reporter construct inhibited expression. Likewise, methylation of all versions of the *TERT* promoter reporters inhibited expression **(Figure 5B)**. Similar observations were made when methylated and unmethylated reporter constructs were transfected in LNCaP and U2OS **(Figure S10)**. Interestingly, reporter constructs driven by a C228T mutant promoter showed approximately 2-fold higher expression than reporter constructs with the WT *TERT* promoter. Methylation of the mutant reporter constructs abolished expression, independent of host cell *TERT* promoter mutational status **(Figure 5C)**. To confirm that methylation of the reporter construct was not dependent on methylation of CpGs outside of the promoter sequence, we used heterologous reporter constructs, pCpGL, in which the backbone plasmid sequence and reporter was engineered without CpGs (36), such that only the TERT promoter would have CpGs. Methylation of these reporters showed uniform repression, independent of host cell. Additionally, we observed that the reporter driven by the C228T mutant promoter showed approximately 3-fold higher expression than the WT reporter construct, independent of host cell *TERT* promoter status **(Figure 5D)**. These functional data suggest that methylation of TERT promoters leads to strong transcriptional silencing regardless of mutation status.

**Figure 5.**
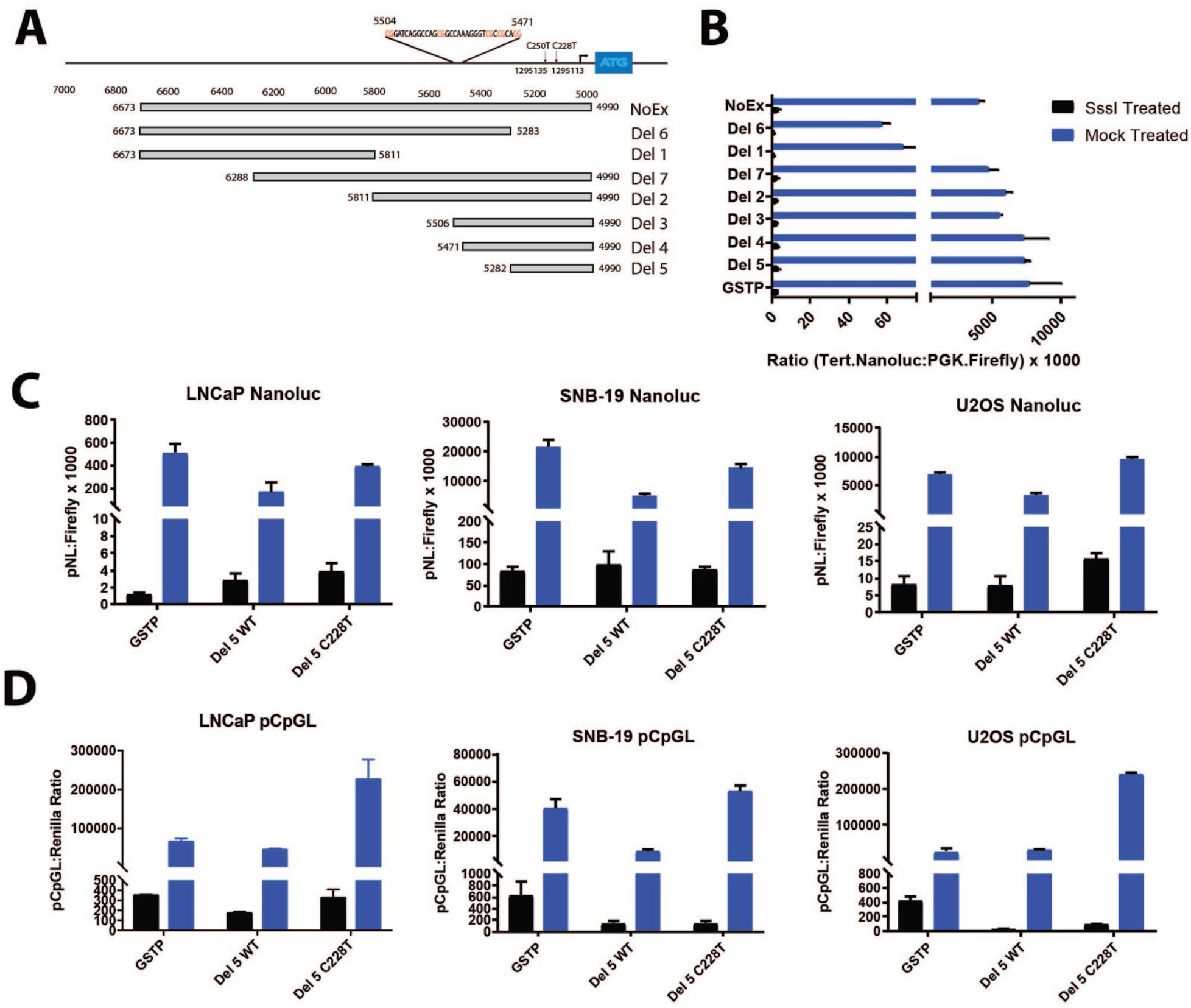
Methylation of *TERT* promoter sequences results in strong repression of heterologous reporter constructs. Reporter assays demonstrated that DNA methylation of the *TERT* promoter greatly suppressed reporter expression in heterologous *TERT* promoter-reporter constructs. (A) Schematic of various deletion constructs of *TERT* promoter driving luciferase expression. (B) HEK293T cells were transfected with deletion constructs of the *TERT* promoter driving luciferase expression with activity similar to a reporter construct driven by the *GSTP1* promoter. The exceptions were deletion constructs lacking elements of the core promoter (Del 6 and Del 1). Methylation of reporters significantly inhibited expression in all constructs. (C) Nanoluc or (D) pCpGL reporters driven by mutant or wild type TERT promoter shows higher expression in mutants compared to wild type promoter, with activity largely abolished in methylated constructs independent of host cell mutation status.

## DISCUSSION

In summary, our findings show that immortalized and cancer cell lines are generally hypermethylated in a region upstream of the recurrent C228T and C250T *TERT* promoter mutations, while normal primary cells are comparatively hypomethylated. These findings are consistent with previous reports describing hypermethylation of the CpG island spanning the *TERT* promoter in cancers compared to normal tissues (25,37–40). However, at the allele level, we find that the hypermethylated alleles are associated with repressed expression, while the remaining unmethylated alleles are associated with active chromatin marks, and are responsible for the observed *TERT* expression in cancer. The association of hypomethylated alleles with active expression and hypermethylated alleles with repressed expression is particularly evident in cancers harboring promoter mutations, and cancers displaying allele specific expression. Consistent with the findings of Stern and colleagues (22), we find that in cancers with *TERT* promoter mutations, the expressed allele is the mutant allele while the silenced allele is WT. Here in this study, we extend these findings by phasing the methylation pattern with the promoter mutation status at a single molecule level and also extent to cancer cell line models with WT *TERT* promoters, wherein the methylated alleles are silenced and the hypomethylated alleles are expressed.

The significance of promoter methylation in the modulation of *TERT* expression is perhaps best represented in the unique cases of cancer cell lines with predominantly mutated *TERT* promoter alleles, such as U251 and HOP62, that possess a balance of both unmethylated and methylated alleles, with open chromatin marks enriched in the unmethylated alleles. These observations suggest that even alleles with *TERT* promoter mutations are silenced by methylation. In our reporter assays of heterologous constructs driven by the *TERT* promoter, we find that introducing the highly recurrent C228T mutation increases reporter expression by approximately 2 to 3-fold, consistent with prior reports on the impact of *TERT* promoter mutations on gene expression (12,41). More importantly, we also find that methylation of these reporter constructs suppresses expression, independent of promoter status as WT or mutant.

Collectively, the findings reported here reaffirm that methylation of *TERT* promoter sequences is a signal of repression rather than activation of expression. The balance of methylated and unmethylated alleles in cancer cells, particularly in the cases where the majority of alleles are methylated and only a fraction of cells have unmethylated and active alleles, suggests that cancer cells behave as a plastic population of stem-like cells. Following telomerase activation, it is intriguing that cancer cells would repress *TERT* expression. One possibility is that repression of all *TERT* alleles by DNA methylation is a default tumor suppressing response in all cells post-telomerase activation, but selective pressure ensures the persistence of unmethylated alleles, as suggested by others (23); after all, cancer cells require some means of telomere maintenance to sustain indefinite proliferation. However, if expression and activation of *TERT* alleles is entirely advantageous to cancer cells, then it is unclear why cells with unmethylated alleles would not, over time, dominate the population, resulting in the majority of alleles being hypomethylated rather than what we observe here.

Another intriguing possibility is that excessive activation of *TERT* may have negative consequences on growth, survival or other cancer phenotypes, and as a result, there is selective advantage for fine tuning the amount of *TERT* activation through DNA methylation-mediated epigenetic repression of some alleles. This would explain the observation that *TERT* expression is relatively low in the majority of cancers, just sufficient to maintain telomere lengths that are already relatively short compared to normal cells (42,43). These findings thus nominate a provocative hypothesis that excessive *TERT* may be disadvantageous in cancer cells, and would be worthwhile to investigate in future studies. In conclusion, our findings suggest that hypermethylation of the *TERT* promoter alleles signals transcriptional repression of those alleles, leading to attenuation of *TERT* activation in cancer cells.

## MATERIALS AND METHODS

### Cell line Validation

A panel of 96 cell culture models were included in this study, 85 cancer lines, 6 immortalized lines, and 5 normal primary cell models. Cell line identify was validated through STR profiling (see Table S1. Cell Line Characteristics).

### Preparation of bisulfite sequencing amplicon libraries and deep sequencing

DNAs were extracted from cells in culture using DNeasy (Qiagen) as per manufacturer’s instructions. DNA (400 ng) from each sample was bisulfite converted using the EZ DNA Methylation-Gold Kit(Zymo) and eluted in 10 μL of water. We used the Accessy Array (Fluidigm, San Francisco, California) platform to generate an amplicon-based library for next generation sequencing, which can process up to 48 samples per chip. To prepare amplicons, 4.88 μL of bisulfite converted DNA was combined with 0.6 μL of 10X JumpStart PCR buffer, 0.12 μL of 10 mM dNTPs, 0.1 μL of JumpStart Taq polymerase (Sigma), and 0.3 μL of Access Array 20X loading buffer. Each sample well of an Access Array chip was loaded with 5 μL of the PCR mixture, and each primer well was loaded with 5 μL of 4 μM primers in 1X loading buffer. Primer sequences can be found in **Table S2**. The following cycling conditions were used: 1 cycle of 50°C for 2 minutes; 1 cycle of 70°C for 20 minutes; 1 cycle of 94°C for 3 minutes; 5 cycles of 94°C for 30 seconds, 57°C for 30 seconds, and 72°C for 90 seconds, with the Tm dropping by 1°C for each cycle; 30 cycles of 94°C for 30 seconds, 51°C for 30 seconds, and 72°C for 90 seconds; and 1 cycle of 72°C for 5 minutes. To barcode and incorporate next generation sequencing adaptors for the Access Array samples, PCR products were diluted 1:100 in water, 1 μL of the diluted PCR product was amplified in a 20 μL barcoding reaction that included 1X NEBNext Phusion Master (NEB) and 4 μL of the Fluidigm Access Array barcoding primers. The following cycling conditions for the barcoding reaction were used: 1 cycle of 98°C for 2 minutes; 15 cycles of 98°C for 30 seconds, 60°C for 30 seconds, 72°C for 90 seconds; and 1 cycle of 72°C for 5 minutes. The barcoded products were then pooled and purified by Ampure XP beads (Beckman Coulter) as per the manufacturer’s instructions. The pooled samples were then subjected to paired end 2×150 bp next generation sequencing using the Fluidigm FL1 (read 1 and read 3 forward and reverse sequencing primers) and FL2 (custom barcode sequencing primer) sequencing primers diluted in HT1 buffer to a final concentration of 500 nM (Fluidigm, see Access Array System for Illumina Sequencing Systems user guide) on a MiSeq or HiSeq 2000 next generation sequencing instrument according to the manufacturer’s protocols (Illumina, San Diego, California). The resulting paired-end reads were demultiplexed using the custom sample barcode sequences to obtain paired end fastq files for each sample. Since all amplicons were less than 300 bp, the 2×150bp paired end reads were used to create consensus merged reads using FLASH (44). These merged reads were then aligned to virtually bisulfite converted reference amplicon sequences using BWA-MEM (45) using the following command to adjust clipping, gap opening, mismatch penalties and bandwidth (bwa mem -B 1 -L 30 -O 30 -w 10). The resulting SAM format alignment files were then parsed using custom scripts to identify positions of mismatches and conversion status of each cytosine in every read from each amplicon and sample. Reads aligning 95% to the converted reference with 5 or fewer mismatches were stored in a final SAM file and summarized in tables. For the Tert_29 amplicon, which overlaps the G->A promoter mutations, additional tables that parsed the mutant and non-mutant reads were generated and the cytosine conversion at each CpG was recorded in tabular format. The resulting tables were analyzed and visualized using R/Bioconductor packages to generate figures. For further interactive analysis/visualization of the data, we used a custom Java program called NextGenDNAMethylMap v1.2 (B.K., S.Y., unpublished).

### Real-Time qPCR to assess *TERT* Expression

RNA was extracted from cells using RNeasy kit (Qiagen). Approximately 2 μg of RNA from each cell line was converted to cDNA using the High-Capacity cDNA Reverse Transcription Kit (Thermo), and subsequently combined with water to make a 1:5 dilution. To measure TERT expression, a 20 μL RT-PCR reaction was prepared with 1X iQ SYBR Green Supermix reagents (Biorad), 3 μL of diluted cDNA, and 500 nM of the primer pair for TERT or TBP (TERT_Ex9-10_forward 5’-AGTGCCAGGGGATCCCGCA, TERT_Ex9-10_reverse 5’-GAGGTGTCACCAACAAGAAATCATCC, TBP_forward 5’-CACGAACCACGGCACTGATT, TBP_reverse 5’-TTTTCTTGCTGCCAGTCTGGAC). The following cycling conditions were used: 1 cycle of 95°C for 3 minutes, 35 cycles of 95°C for 25 seconds and 60°C for 30 seconds. The expression of *TERT* was normalized to *TBP* expression (2^−ΔCt^).

### *TERT* promoter bisulfite Sanger sequencing

DNA from cells were extracted using DNeasy (QIAgen). DNA (100 ng) was bisulfite converted using the EZ DNA Methylation Gold Kit (Zymo). Bisulfite converted DNA was eluted in 30 μl water, of which 10 μl was used for a 40 μL PCR reaction containing 1X Buffer, 1.5 mM MgCl_2_, 250 nM each dNTPs, 12.5 μg BSA, 6.25% DMSO, 5 units of Platinum Taq (Lifetech) and 400 nM of forward and reverse primers (TertMut_BSF2_forward 5’-GAAAGGAAGGGGAGGGGTTGGGAGGG and TertMut_BSF2_reverse 5’-CCTCCACATCATAACCCCTCCCT) primers, and 5 units of Platinum Taq (Lifetech). The following cycling conditions were used: 1 cycle of 95°C for 3 min; 35 cycles of 95°C for 30 seconds, 56°C for 30 seconds, and 72°C for 1 minute; and 1 cycle of 72°C for 5 minutes. Products were run on a 2% agarose gel, excised, and cloned into the pCR2.1-TOPO vector system (Lifetech). Vector was transformed in TOP10 chemically competent cells (Lifetech). Plasmid DNA was purified from colonies using the Plasmid Purification Kit (Qiagen), and subsequently analyzed by Sanger sequencing. Sanger Bisulfite Sequencing data were analyzed using a custom Java program called DNAMethylMap which facilitates analysis of Sanger bisulfite sequencing clones with virtually bisulfite converted reference amplicon sequences (B.K., S.Y., unpublished).

### Long-range bisulfite sequencing

To interrogate a longer amplicon (1651 bp), we modeled our approach after single-molecule real-time bisulfite sequencing (SMRT-BS) (46). Genomic DNA (2.4 μg) extracted from cells were bisulfite converted using the Methylamp DNA Modification Kit (Epigentek). The region of interest was PCR amplified in four 50 μL reactions using the following conditions: 1X JumpStart Buffer, 1M Betaine (Sigma), 2.5 μL of JumpStart Taq DNA Polymerase (Sigma) per 50 L reaction, 500 nM of primers (TERT_Long_forward 5’-GGATTTGGAGGTAGTTTTGGGTTTT and TERT_Long_reverse 5’-CCTAAAAAATAAAAAAAATACTTAATCTC). The following cycling conditions were used: 1 cycle of 94°C for 105 seconds; 40 cycles of 94°C for 30 seconds, 50°C for 30 seconds, and 65°C for 5 minutes; and 1 cycle of 65°C for 5 minutes. PCR products were dA tailed (NEBNext® dA-Tailing Module), cloned into TOPO TA vectors, and individual clones were sanger sequenced and analyzed with DNAMethylMap software (B.K., S.Y., unpublished).

### Sanger sequencing of rs2736098 for genomic DNA and cDNA

Genomic DNA was extracted from cells in culture using DNeasy (Qiagen). RNA was extracted from cells using the RNeasy kit (Qiagen), and subsequently converted into cDNA using the High-Capacity cDNA Reverse Transcription Kit (Thermofisher). Genomic DNA or cDNA (~30 ng) PCR amplified in a 40 μL PCR reaction containing 1X Buffer, 1.5 mM MgCl_2_, 250 μM each dNTPs, 10 μg BSA, 5% DMSO, 3 units of Platinum Taq (Lifetech) and 400 nM of primers (rs2736098_forward 5’-CCTTGTCGCCTGAGGAGTAG and rs2736098_reverse 5’-GTGACCGTGGTTTCTGTGTG). The following cycling conditions were used: 1 cycle of 95°C for 3 minutes; 32 cycles of 95°C for 30 seconds, 56°C for 30 seconds, and 72°C for 40 seconds; and 1 cycle of 72°C for 5 minutes. PCR products were purified using DNA Clean and Concentrator-5 columns (Zymo) and subsequently analyzed by Sanger sequencing.

### Preparation of luciferase constructs for methylated/unmethylated promoter bashing

*TERT* promoter regions were cloned into the pNL1.1 vector (Promega), transformed in TOP10 chemically competent cells (Lifetech), and verified by Sanger sequencing. To generate methylated versus unmethylated plasmids, 1 μg of plasmid was treated with SssI (NEB) or mock as per the manufacturer’s instructions. To ensure complete methylation, plasmid DNA was treated with SssI twice. Cells in culture were seeded into 96 well plates and transfected in suspension using X-tremeGENE HP DNA Transfection Reagent (Roche). Transfection complexes were prepared such that each reaction contained 11 μL of Opti-MEM, 59 ng of empty pCR2.1-TOPO vector, 5 ng of pGL4.53(luc2/PGK) vector (Promega), 1 ng of pNL plasmid, and 0.12 μL of HP transfection reagent. After two days, transfected cells were lysed in 50 uL of passive lysis buffer (Promega), transferred to black 96 well plates, and measured for reporter activity by mixing 50 μl of ONE-Glo EX Luciferase Reagent and NanoDLR Stop & Glo Reagent sequentially as described in the Nano-Glo Dual Luciferase Assay System manual (Promega).

### Preparation of linearized luciferase constructs containing the C228T mutation

We designed a C228T mutant version of the Del 5 *TERT* promoter construct. To ensure that our constructs were interrogating only promoter methylation (*TERT* or *GSTP1*) and to improve the efficiency of transfection, we generated linearized reporter constructs. The promoter region and nanoluciferase coding region was PCR amplified from the pNL1.1 reporter constructs using Phusion High-Fidelity PCR Master Mix with GC Buffer (NEB), and 400 nM of forward and reverse primers (pNL1.1_forward 5’-AATTATCTTAAGATTTCTCTGGCCTAACTGGCCGG and pNL1.1_reverse 5’-AATTATCTTAAGTGGGTTGAAGGCTCTCAAGGGCATC). PCR products were subsequently digested with DpnI (NEB) to eliminate residual plasmid sequence and purified with QIAquick PCR Purification columns (Qiagen). The linearized constructs were subjected to SssI methylation (or mock reactions) as described above, and transfected into cultured cells in 96-well plates. Each well was treated with a transfection mixture containing 11.6 μL of Opti-MEM, 48 ng of pUC18 DNA, 1 ng of linearized nanoluciferase construct, 10 ng of pGL4.53(luc2/PGK), and 0.12 μL X-tremeGENE HP. After two days, transfected cells were analyzed for reporter activity using the Nano-Glo Dual Luciferase Assay System (Promega) as described above.

### Preparation of pCpGL constructs for C228T/wild-type promoter analysis

To confirm that methylation of non-promoter sequences did not significantly affect reporter activity, we obtained the pCpGL plasmid as the kind gift of Dr. Shaohui Hu from the Heng Zu Lab at JHMI. The pCpGL reporter plasmid lacks CpG dinucleotides, therefore, only promoter sequences will contain CpG residues. The *TERT* Del 5, *TERT* Del 5 with C228T, and GSTP1 promoters were cloned into pCpGL to generate 3 unique constructs. The reporter plasmids were treated with SssI or mock, and transfected into cells cultured in 96-well plates. Each well was treated with a transfection mixture containing 11.6 μL of Opti-MEM, 54 ng of pCpGL vector, 4 ng of Renilla (pRL-CMV, Promega), and 0.12 μL of X-tremeGENE HP. After two days, transfected cells were analyzed for reporter activity using the Nano-Glo Dual Luciferase Assay System (Promega) as described above.

### Chromatin immunoprecipitation (ChIP) for open chromatin marks, H3K4me^3^ and H3K27ac, at WT and mutant *TERT* promoter alleles

Adherent cells were fixed for 10 minutes at room temperature (RT) in 1% formaldehyde in PBS. Glycine was added to a final concentration of 125 mM and incubated at RT for 5 minutes. Cells were washed with cold PBS, scraped, pelleted by centrifugation, the supernatant removed, resuspended in 5 mL of LB1 (50 mM HEPES, pH 8, 140 mM NaCl, 1 mM EDTA, 10% glycerol, 0.5% Igepal, 0.25% Triton X-100), and incubated on ice for fifteen minutes. Cells were subsequently pelleted, the supernatant removed, resuspended in 5 mL of LB2 (10 mM Tris-HCl, pH 8, 200 mM NaCl, 1 mM EDTA, 0.5 mM EGTA) and incubated for 10 minutes on ice. Cells were pelleted, the supernatant removed, and resuspended in 1 mL of Shearing Buffer (0.2% SDS, 50 mM Tris-HCl, pH 8, 5 mM EDTA, 1X cOmplete EDTA-free protease inhibitor cocktail). Cells were sonicated using the Covaris S2 and centrifuged at 16,000 xg for 10 minutes at 4°C. Lysate supernatant (1 mL) was mixed with II mL of dilution buffer (2 mM EDTA, 150 mM NaCl, 20 mM Tris-HCl, pH 8, 1X complete EDTA-free protease inhibitor cocktail). Diluted lysate (1.9 mL) was incubated overnight at 4°C with one of the following antibodies: 31 μL H3K4me^3^ (ab8580, abcam), 5 μL H3K27ac (ab4729 abcam), or 4 μL IgG (2729, Cell Signaling). Dynabead Protein G beads (30 μL) were combined with IP mixtures, and incubated at 4°C for 3 hours. Beads were precipitated and washed sequentially with TSE I (0.1% SDS, 1% Triton, 2 mM EDTA, 20 mM Tris-HCl, pH 8, 150 mM NaCl), TSE II (0.1% SDS, 1% Triton, 2 mM EDTA, 20 mM Tris-HCl, pH 8, 300 mM NaCl), TSE III (0.25 M LiCl, 1% Igepal, 1 mM EDTA, 10 mM Tris-HCl, pH 8, 1% deoxycholate), and finally TE. Washed beads were incubated with 100 μL of Elution Buffer (1% SDS, 0.75% sodium bicarbonate) at 55°C for 15 minutes. Crosslinks in samples were reversed by incubated at 65°C for 8 hours, and purified using the QIAquick PCR Purification Kit (Qiagen) and eluted in 100 μL water. TERT promoter bisulfite Sanger sequencing was performed as described above.

### *TERT* promoter ChIP and bisulfite sequencing

The iDeal ChIP-Seq Kit for Transcription Factors (Diagenode) was used to prepare ChIP libraries according to manufacturer’s instructions. Briefly, cells were fixed for 10 minutes at RT in 11% formaldehyde solution in fixation buffer (Diagenode), diluted 1:10 in RPMI with 10% FBS. The fixation reaction was quenched with glycine solution (Diagenode) and incubated at room temperature for 5 minutes. Fixed cells were washed and resuspended in sonication buffer supplemented with protease inhibitors (from Diagenode) and PhosStop (Roche), and subsequently sonicated using the Covaris S2 sonicator. ChIP for RNA polymerase II was performed according the iDeal ChIP-Seq Kit instructions using the anti-Pol2 antibody (Abcam, ab5408). Immunoprecipitated DNA was bisulfite converted using the EZ DNA Methylation Gold Kit (Zymo), and PCR amplified. Bisulfite converted DNA was put into a 40 uL PCR reaction comprised of 1X Buffer, 1.5 mM MgCl_2_, 250 nM of dNTPs, 1 M betaine, 0.5 uL of Platinum Taq (Lifetech) and 400 nM of primers (mini_BSF_forward 5’-GTTGGAAGGTGAAGGGGTAGG, mini_BSF_reverse 5’-TCCCTACACCCTAAAAAC). The following reaction conditions were used: 1 cycle of 95°C for 3 min; 40 cycles of 95°C for 30 seconds, 53°C for 30 seconds, and 72°C for 45 seconds; and 1 cycle of 72°C for 5 minutes. PCR products were cloned, transformed and Sanger sequenced as described above.

### Real-Time qPCR of ChIP libraries

Each PCR were carried out in 20 uL reactions containing 3 uL of ChIP library, 1X SsoAdvanced Universal SYBR Green Supermix (Biorad), and 500 nM primers. Due to the GC-rich content of the TERT promoter region, 1M Resolution Solution (GC-RICH PCR, Roche) was added to each qPCR reaction (47). The following primer sets were used: Tert_ Mut_3_forward 5’-CGCGCGGACCCCGCCCCGTCCCGAC and Tert_ Mut_3_reverse 5’-ACGCAGCGCTGCCTGAAACTCGCGC, ACTB_forward 5’-AAGGCGAGGCTCTGTGCT and ACTB_reverse 5’-CCGAAAGTTGCCTTTTATGG, PSA7_forward 5’-TGGGACAACTTGCAAACCTG and PSA7_reverse 5’-CCAGAGTAGGTCTGTTTTCAATCCA. The following thermocycling conditions were used: 1 cycle of 95°C for 3 minutes, 35 cycles of 95°C for 25 seconds and 60°C for 30 seconds.

## ACKNOWLEDGMENTS

The pNL1.1 vector was the kind gift of May Guo (Promega). We thank Dr. Theodore L. DeWeese for helpful discussions. We thank the members of the SKCCC Experimental and Computational Genomics Core, supported by the Johns Hopkins Sidney Kimmel Comprehensive Cancer Center Regional Oncology Center Grant (NIH/NCI P30CA006973). This work was supported in part by grants from the NIH/NCI (P50CA058236, U01CA196390, R01CA183965), Commonwealth Foundation, and the Prostate Cancer Foundation, the V Foundation For Cancer Research, the Masenheimer Fund, the Irving Hansen Foundation, and the Patrick C. Walsh Award.

