## Supplemental Materials for "Pervasive promoter hypermethylation of silenced *TERT* alleles in human cancers"

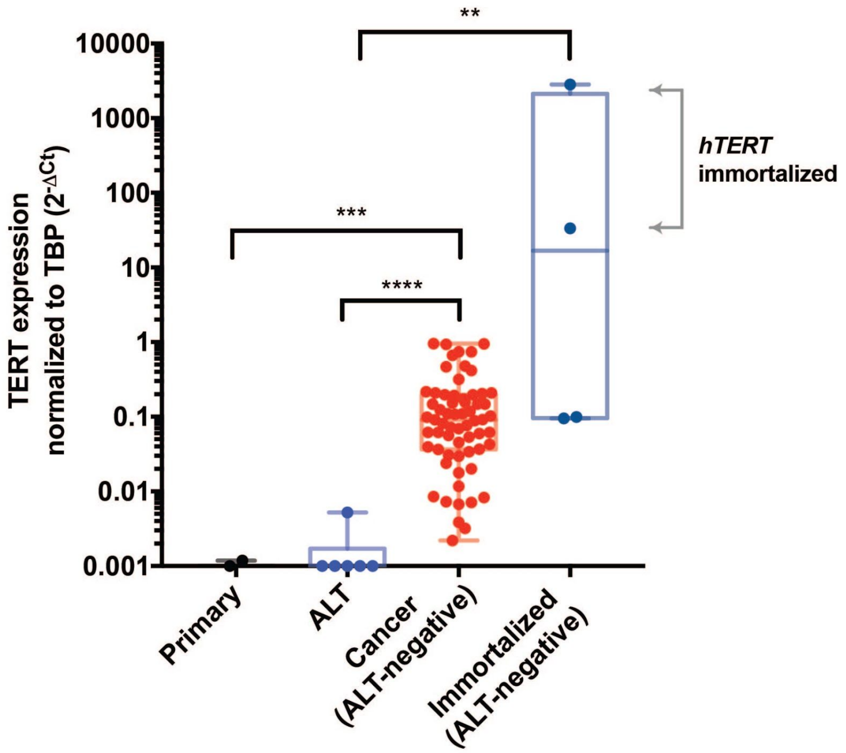


**Figure S1.** ***TERT*** **is expressed in ALT-negative cancer cell lines and TERT-immortalized primary cells.** Mann-Whitney test was used to assess statistically significant difference. Asterisks denote level of significance (* $p \leq 0.5$, ** $p \leq 0.01$, *** $p \leq 0.001$, **** $p \leq 0.0001$).

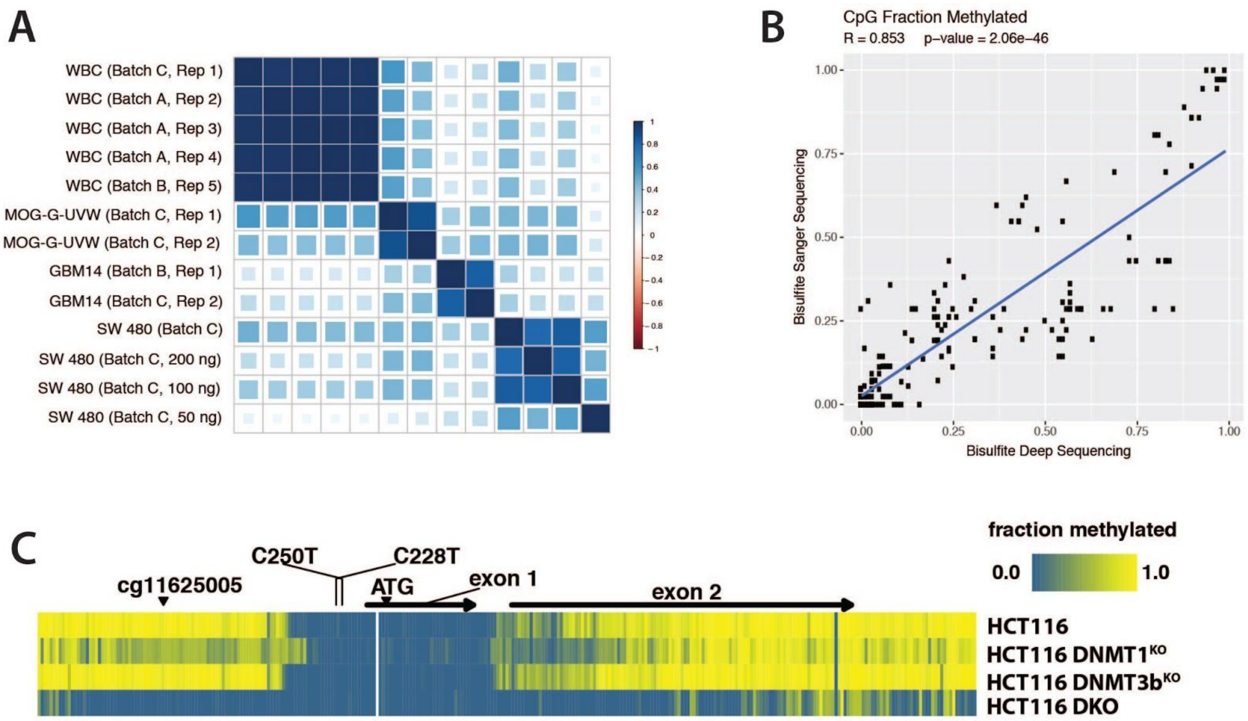


**Figure S2. Ultra-deep bisulfite sequencing measurements of the *TERT* promoter region are highly robust.** (A) Correlation measurements show high reproducibility and precision in replicate samples including intra- and inter-batch controls. (B) There is high concordance of ultra-deep bisulfite sequencing measurements with bisulfite sanger sequencing. (C) As expected, DNA methyltransferase (DNMT) deficient cells (HCT116 DKO) are unmethylated compared to WT parental HCT116 cells.

A

B

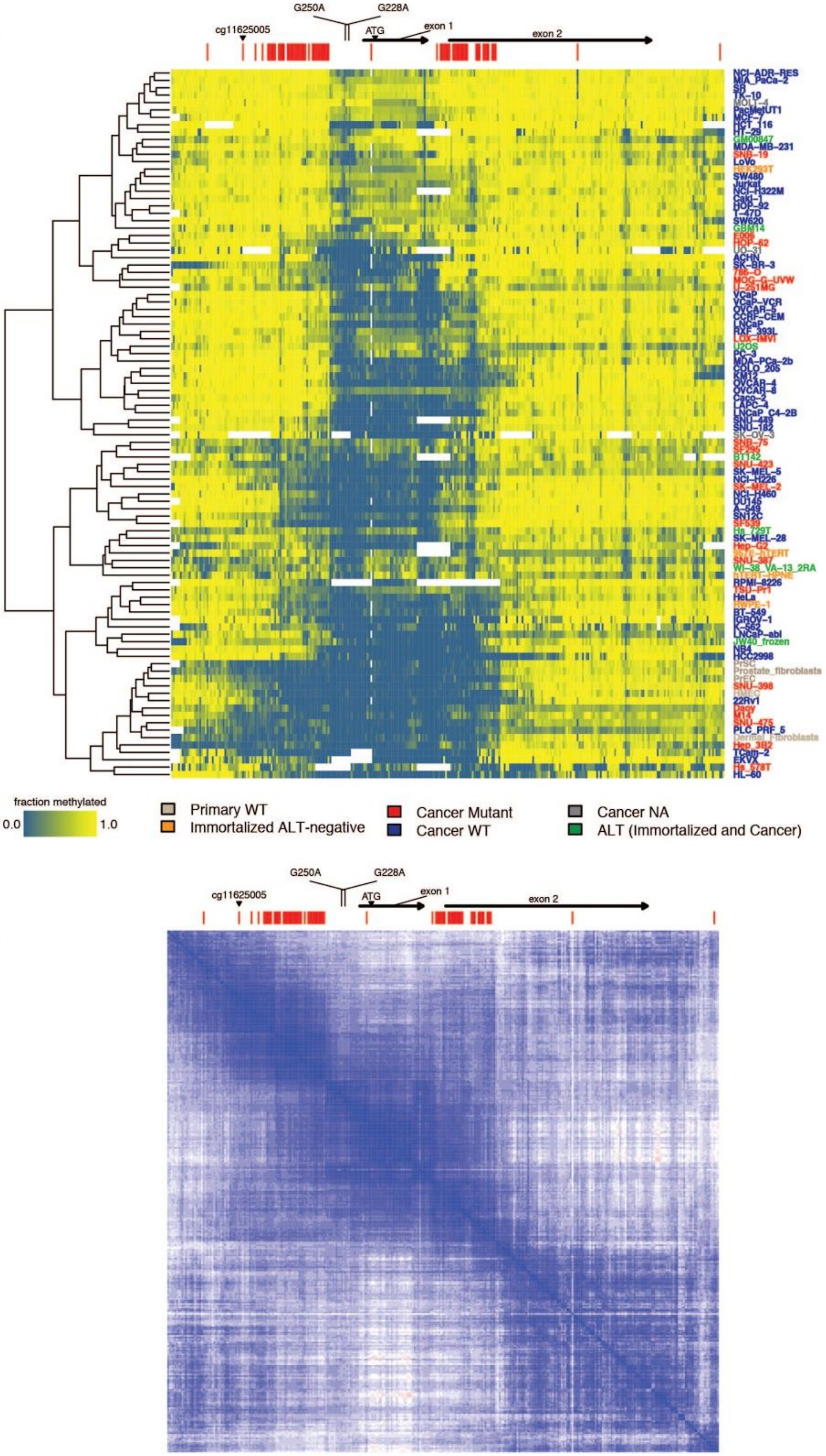

G250A
G228A
cg11625005
ATG
exon 1
exon 2
NCI-ADR-RES
MIA_PaCa-2
SR
TK-10
MOLT-4
PacMetUT1
MCF-7
HCT_116
HT-29
GM00847
MDA-MB-231
SNB-19
LoVo
HEK293T
SW480
Jurkat
NCI-H322M
Caki-1
HOP-92
T-47D
SW620
GBM14
E006
HOP-62
UO-31
ACHN
SK-BR-3
786-O
MOG-G-UVW
U-251MG
VCaP
VCaP-VCR
OVCAR-5
CCRF-CEM
LNCaP
RXF_393L
LOX-IMVI
U2OS
PC-3
MDA-PCa-2b
COLO_205
KM12
OVCAR-4
OVCAR-8
Caco-2
LAPC-4
LNCaP_C4-2B
SNU-449
SNU-182
SK-OV-3
SNB-75
SF295
BT142
SNU-423
SK-MEL-5
NCI-H226
SK-MEL-2
NCI-H460
DU145
A-549
SN12C
SF539
Hs_729T
SK-MEL-28
Hep-G2
SNU-387
WI-38_VA-13_2RA
hTERT-HPNE
RPMI-8226
TSU-Pr1
HeLa
RWPE-1
BT-549
IGROV-1
K-562
LNCaP-abl
JW40_frozen
NB4
HCC2998
PrSC
Prostate_fibroblasts
PrEC
SNU-398
HMEC
22Rv1
Daoy
M14
SNU-475
PLC_PRF_5
Dermal_Fibroblasts
Hep_3B2
TCam-2
EKVX
Hs_578T
HL-60
fraction methylated
0.0
1.0
Primary WT
Immortalized ALT-negative
Cancer Mutant
Cancer WT
Cancer NA
ALT (Immortalized and Cancer)

**Figure S3. CpG methylation measurements of the *TERT* promoter.** (A) Clustering samples according to top 20% most variable CpGs, normal primary samples are generally the most hypomethylated. (B) A correlation matrix of all CpGs by all CpGs ordered by position show blocks of highly correlated CpGs on either side of the highly recurrent promoter mutations. Red hashes show the position of the top 20% most variable CpGs.

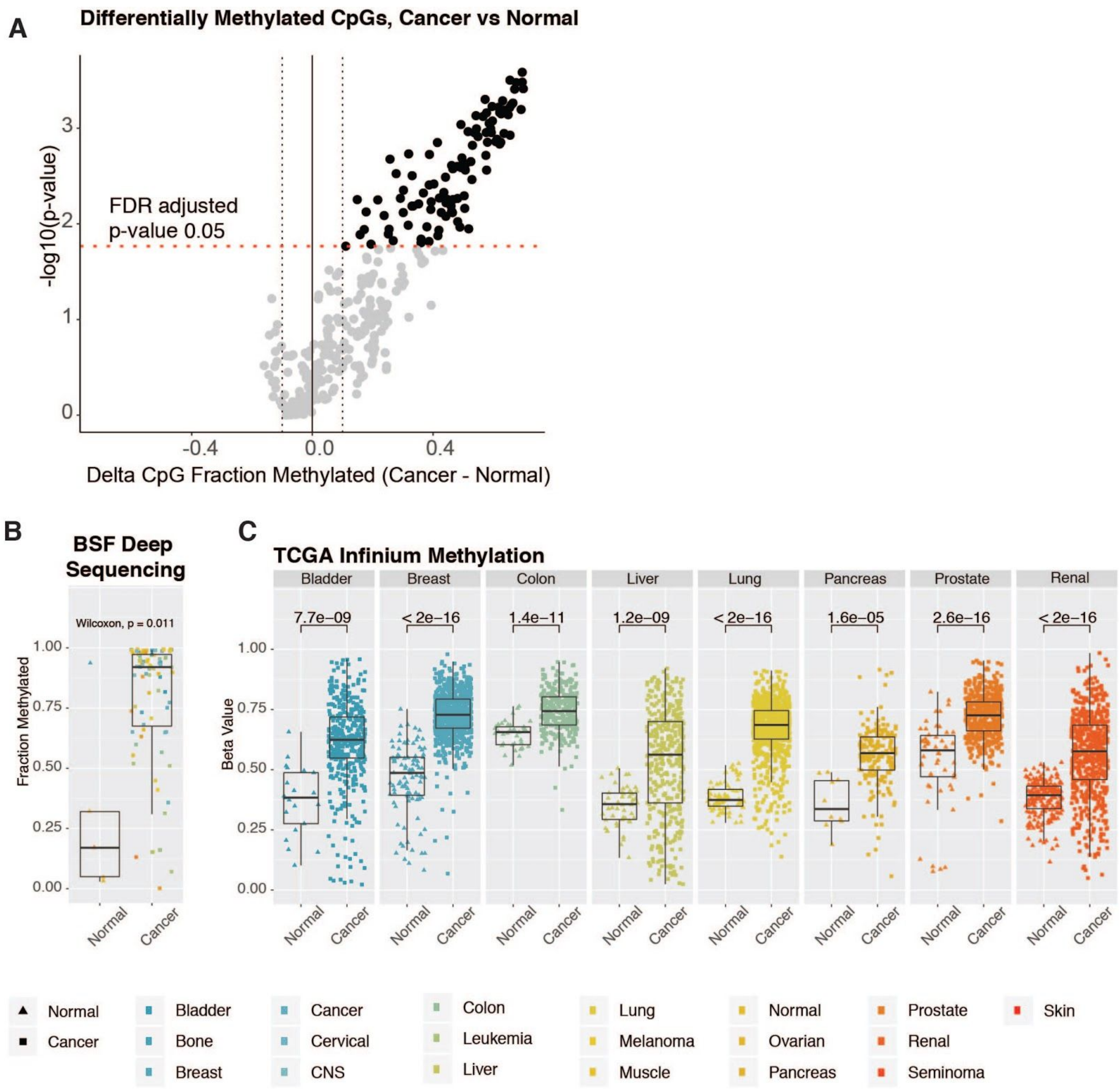


**Figure S4. The *TERT* promoter region is generally hypermethylated in cancers compared to normal.** In ultra-deep BSF sequencing results, (A) the differentially methylated CpGs (delta CpG > 0.1 or delta CpG < -0.1, and FDR adjusted p-value < 0.05) were all more methylated in cancer compared to normal, and (B) cancers were typically more methylated at the CpG position associated with Infinium probe cg11625005. (C) Analysis of Infinium probe cg11625005 in publically available data (TCGA data from cancers also covered by our cell lines) show that cancers are typically more methylated than normal.

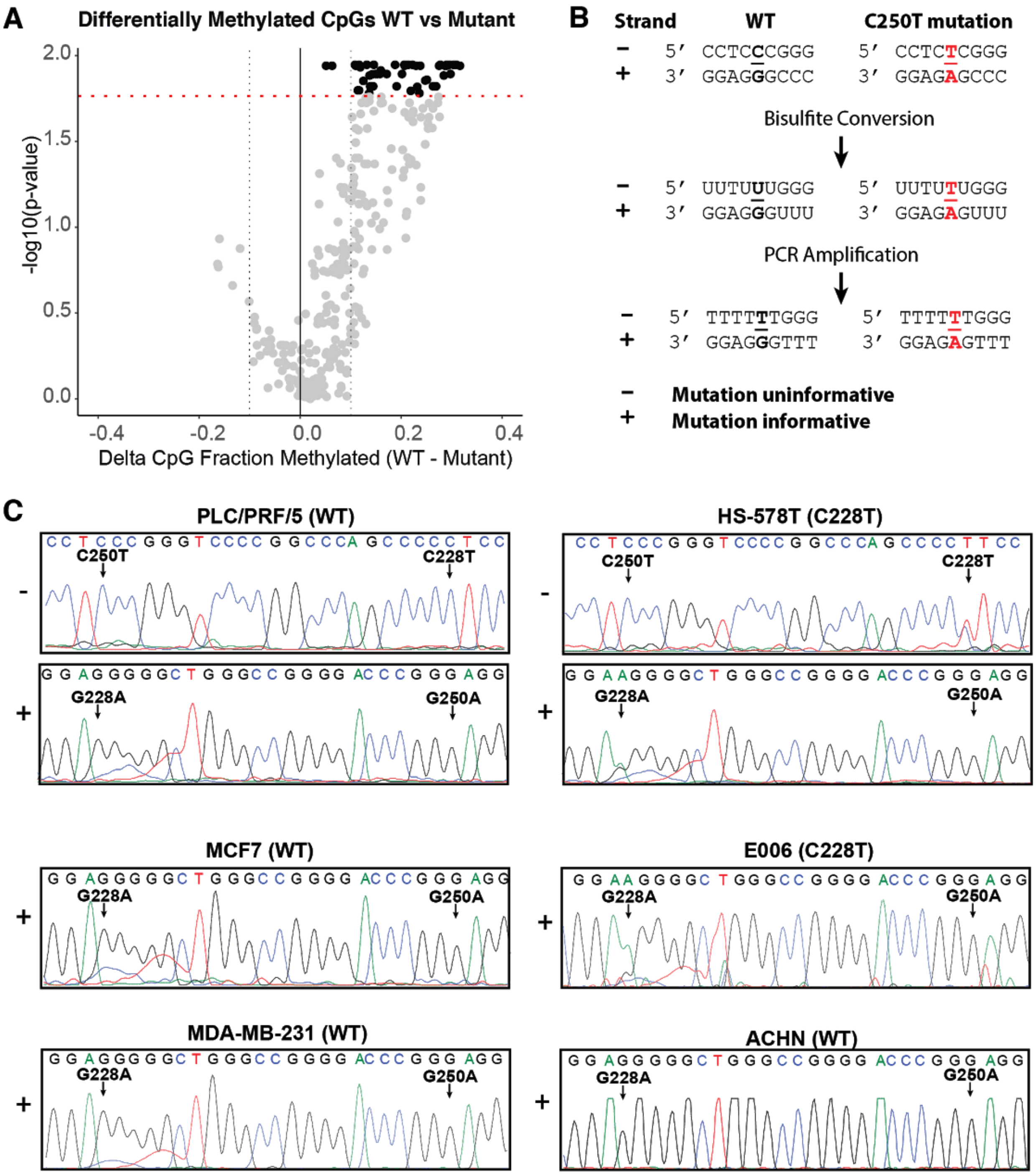


**Figure S5. The *TERT* promoter region is generally hypermethylated in cancers with WT *TERT* promoter compared to cancers with *TERT* promoter mutations.** (A) In ultra-deep BSF sequencing results, the differentially methylated CpGs (delta CpG > 0.1 or delta CpG < -0.1, and FDR adjusted p-value < 0.05) were all more methylated in cancers with WT *TERT* promoters compared to cancers with *TERT* promoter mutations. (B) C -> T mutations can confounded by bisulfite conversion, making them difficult to assess in bisulfite sequencing data. In the C-rich negative strand, the sequencing data is identical between WT and mutant *TERT* promoter sequences following bisulfite conversion and PCR amplification. In contrast, the G-rich positive strand can be used to distinguish WT and mutant sequences following bisulfite conversion and PCR amplification. (C) Sanger sequencing of a subset of the cancer cell lines confirm deep sequencing results of WT and mutant *TERT* promoter. The plus and minus strands of PLC/PRF/5 and HS-578T were sequenced by Sanger. The minus strand of MCF7, MDA-MB-231, E006 and ACHN were sequenced by Sanger.

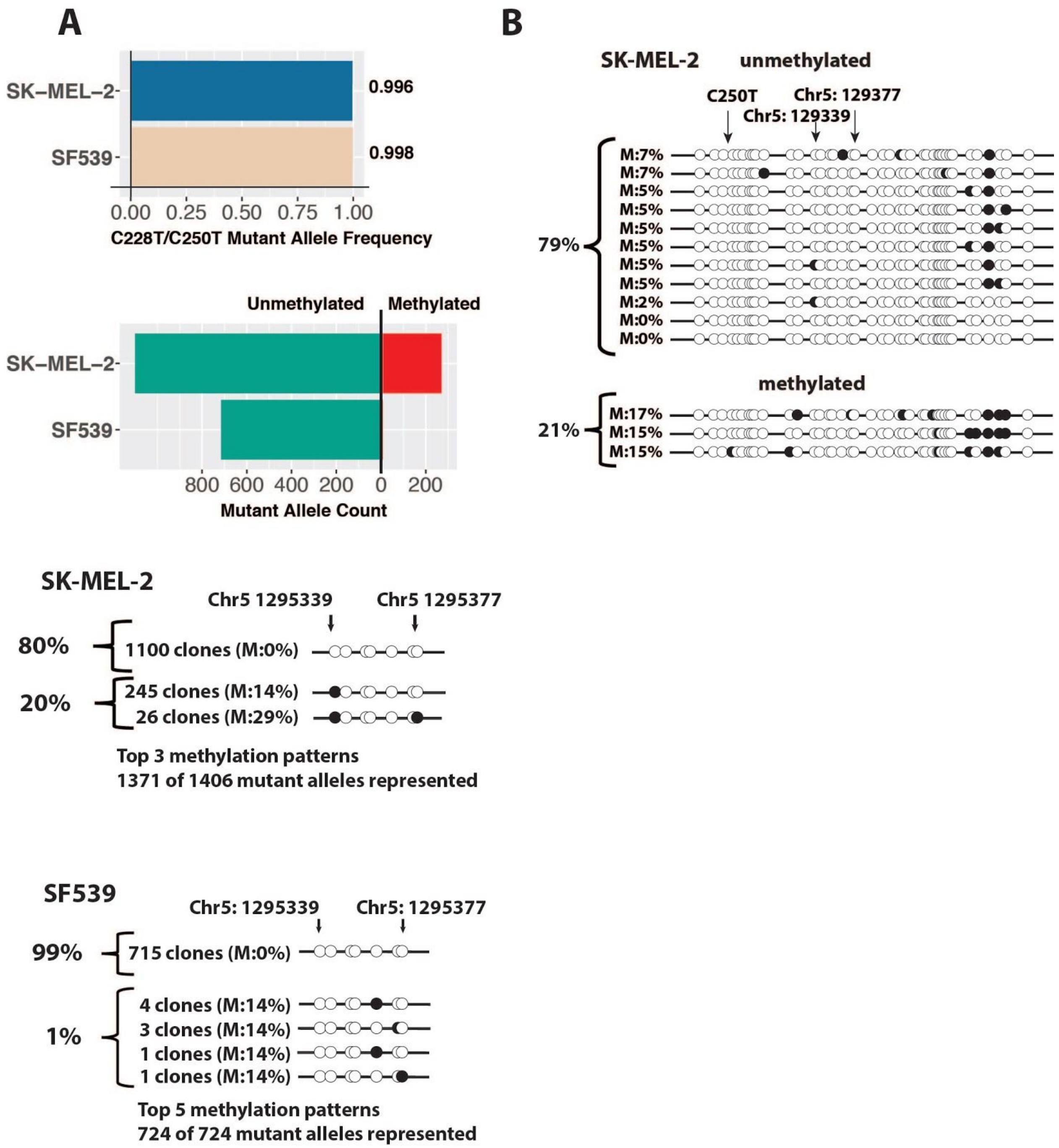


**Figure S6. Cancer cells with high mutant allele fractions with predominantly unmethylated active alleles.** (A) BSF deep sequencing indicate that more than 99% of the alleles of SK-MEL-2 and SF539 have TERT promoter mutations, and 80% of the alleles are unmethylated, consistent with previous observations that mutant alleles are generally hypomethylated. Black circles indicate methylated CpGs and open circles indicate unmethylated

CpGs. (B) Bisulfite Sanger sequencing of SK-MEL-2 confirm that the majority of the mutant alleles are unmethylated.

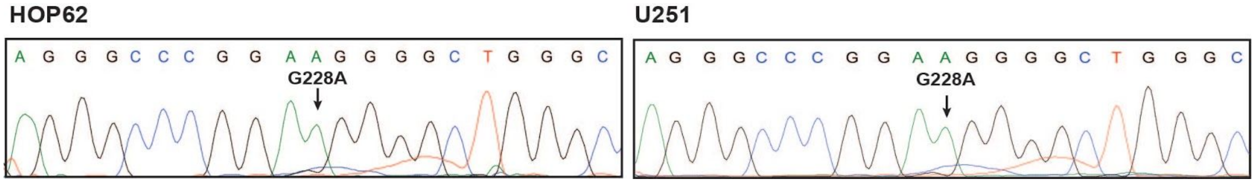


**Figure S7. U251 and HOP62 are homozygous for *TERT* promoter mutations**. Sanger sequencing of genomic DNA confirmed that U251 and HOP62 alleles have TERT promoter mutations on all alleles.

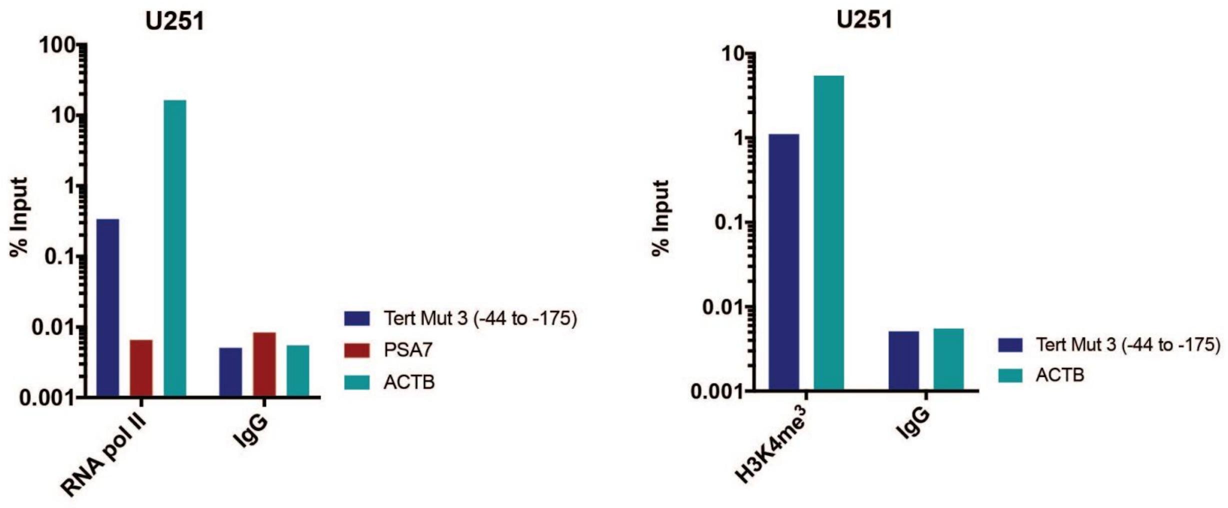


**Figure S8. Active chromatin marks bound to *TERT* promoter mutant sequence.** ChIP-qPCR data for IgG, H3K4me3 and Pol2 in U251 cells showed enriched binding to the *TERT* mutant. *ACTB* is a highly expressed gene, and is expected to be enriched for Pol2 and H3K4me3. *PSA7* is used as a negative control in these ChIP-qPCR assessments.

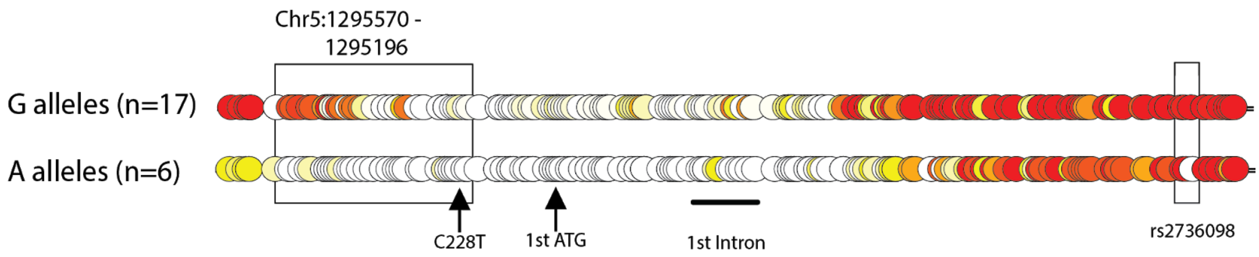


**Figure S9. Long-range bisulfite amplicon sequencing of *TERT* promoter in RPMI-8226.** Methylation map of BSF sequencing encompassed the core TERT promoter, including the region upstream of the highly recurrent promoter mutations, and the heterozygous rs2736098 SNP located in exon 2 of TERT (~1.5 kb).

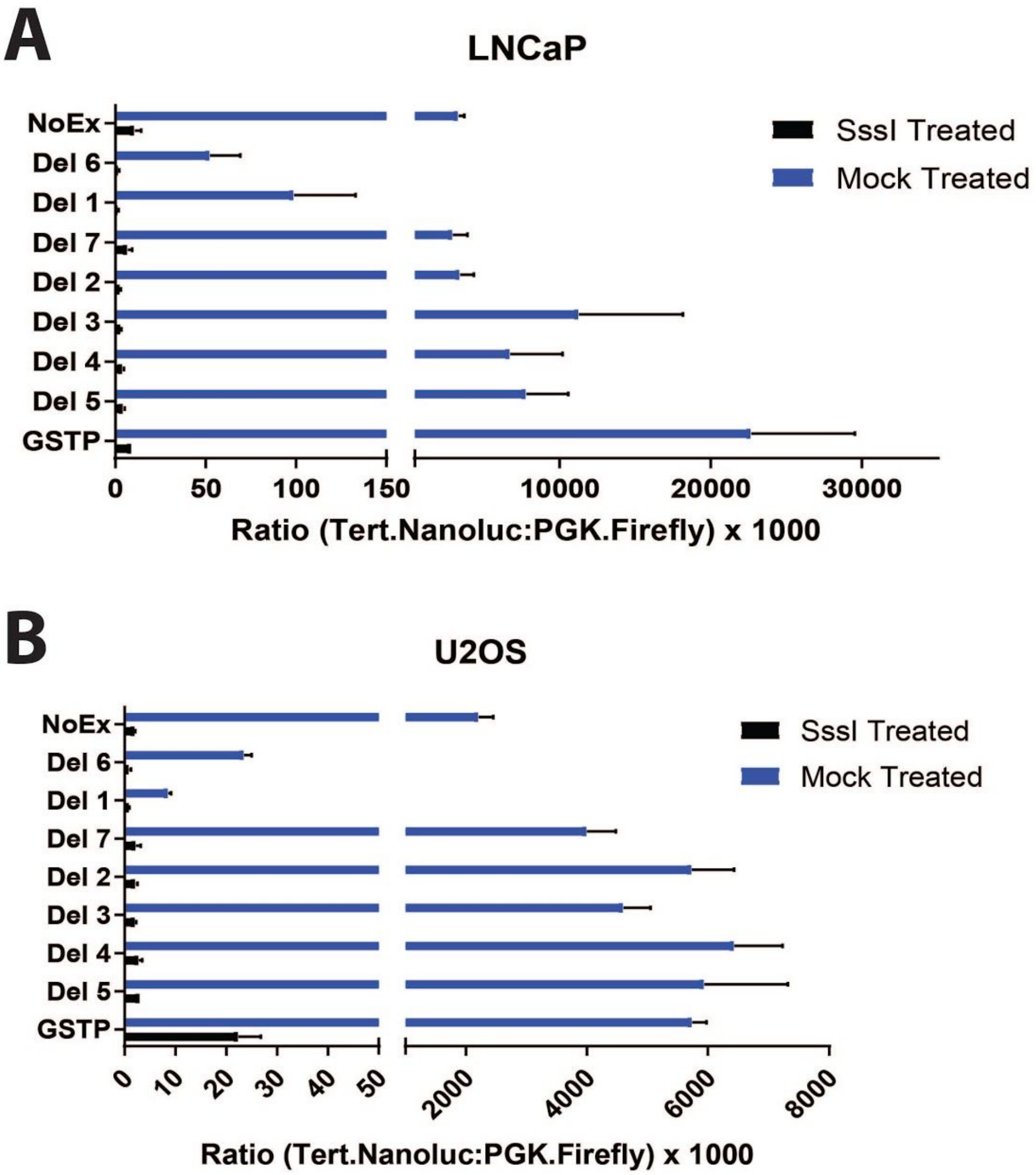


**Figure S10**. **Methylation of *TERT* promoter sequences results in strong repression of heterologous reporter constructs transfected in LNCaP and U2OS.** Cells transfected with various deletion constructs of the TERT promoter driving luciferase expression show activity similar to GSTP1, except for deletions lacking elements of the core promoter. Methylation of reporters significantly inhibited expression of all versions of the reporter constructs.

### TABLES

#### Table S1. Cell line characteristics.

| Cell Line | Tissue Type | Cell Type [a] | TERT Promoter [b] | ALT Status [c] | STR Status [d] | Cell Line | Tissue Type | Cell Type [a] | TERT Promoter [b] | ALT Status [c] | STR Status [d] |
|---|---|---|---|---|---|---|---|---|---|---|---|
| Dermal_Fibroblasts | Skin | Primary | WT | Negative | Pass | 22Rv1 | Prostate | Cancer | WT | Negative | Pass |
| HMEC | Breast | Primary | WT | Negative | Pass | A-549 | Lung | Cancer | WT | Negative | Pass |
| PrEC | Prostate | Primary | WT | Negative | Pass | ACHN | Renal | Cancer | WT | Negative | Pass |
| Prostate_fibroblasts | Prostate | Primary | WT | Negative | Pass | BT-549 | Breast | Cancer | WT | Negative | Pass |
| PrSC | Prostate | Primary | WT | Negative | Pass | CCRF-CEM | Leukemia | Cancer | WT | Negative | Pass |
| RWPE-1 | Prostate | Imm_HPV | WT | Negative | Pass | COLO_205 | Colon | Cancer | WT | Negative | Pass |
| GM00847 | Fibroblast | Imm_SV40 | WT | ALT | Pass | DU145 | Prostate | Cancer | WT | Negative | Pass |
| WI-38_VA-13_2RA | Lung | Imm_SV40 | WT | ALT | Pass | EKVX | Lung | Cancer | WT | Negative | Pass |
| HEK293T | Renal | Imm_SV40 | WT | Negative | Pass | HCC2998 | Colon | Cancer | WT | Negative | Pass |
| 957E-hTERT | Prostate | Imm_TERT | WT | Negative | Pass | HCT_116 | Colon | Cancer | WT | Negative | Pass |
| hTERT-HPNE | Pancreas | Imm_TERT | WT | Negative | Pass | HL-60 | Leukemia | Cancer | WT | Negative | Pass |
| TSU-Pr1 | Bladder | Cancer | Mut | NA | NA | HOP-92 | Lung | Cancer | WT | Negative | Pass |
| 786-O | Renal | Cancer | Mut | Negative | Pass | HT-29 | Colon | Cancer | WT | Negative | Pass |
| Daoy | CNS | Cancer | Mut | Negative | Pass | IGROV-1 | Ovarian | Cancer | WT | Negative | Pass |
| E006 | Prostate | Cancer | Mut | Negative | Pass | K-562 | Leukemia | Cancer | WT | Negative | Pass |
| Hep_3B2 | Liver | Cancer | Mut | Negative | Pass | KM12 | Colon | Cancer | WT | Negative | Pass |
| Hep-G2 | Liver | Cancer | Mut | Negative | Pass | LAPC-4 | Prostate | Cancer | WT | Negative | Pass |
| HOP-62 | Lung | Cancer | Mut | Negative | Pass | LNCaP | Prostate | Cancer | WT | Negative | Pass |
| Hs_578T | Breast | Cancer | Mut | Negative | Pass | LNCaP_C4-2B | Prostate | Cancer | WT | Negative | Pass |
| LOX-IMVI | Melanoma | Cancer | Mut | Negative | Pass | LNCaP-abl | Prostate | Cancer | WT | Negative | Pass |
| M14 | Melanoma | Cancer | Mut | Negative | Pass | MCF-7 | Breast | Cancer | WT | Negative | Pass |
| MOG-G-UVW | CNS | Cancer | Mut | Negative | Pass | MDA-MB-231 | Breast | Cancer | WT | Negative | Pass |
| SF295 | CNS | Cancer | Mut | Negative | Pass | MDA-PCa-2b | Prostate | Cancer | WT | Negative | Pass |
| SF539 | CNS | Cancer | Mut | Negative | Pass | NCI-ADR-RES | Ovarian | Cancer | WT | Negative | Pass |
| SK-MEL-2 | Melanoma | Cancer | Mut | Negative | Pass | NCI-H226 | Lung | Cancer | WT | Negative | Pass |
| SNB-19 | CNS | Cancer | Mut | Negative | Pass | NCI-H322M | Lung | Cancer | WT | Negative | Pass |
| SNB-75 | CNS | Cancer | Mut | Negative | Pass | NCI-H460 | Lung | Cancer | WT | Negative | Pass |
| SNU-387 | Liver | Cancer | Mut | Negative | Pass | OVCAR-4 | Ovarian | Cancer | WT | Negative | Pass |
| SNU-398 | Liver | Cancer | Mut | Negative | Pass | OVCAR-5 | Ovarian | Cancer | WT | Negative | Pass |
| SNU-423 | Liver | Cancer | Mut | Negative | Pass | OVCAR-8 | Ovarian | Cancer | WT | Negative | Pass |
| SNU-475 | Liver | Cancer | Mut | Negative | Pass | PacMetUT1 | Prostate | Cancer | WT | Negative | Pass |
| U-251MG | CNS | Cancer | Mut | Negative | Pass | PC-3 | Prostate | Cancer | WT | Negative | Pass |
| MOLT-4 | Leukemia | Cancer | NA | Negative | Pass | PLC_PRF_5 | Liver | Cancer | WT | Negative | Pass |
| SK-OV-3 | Ovarian | Cancer | NA | Negative | Pass | RPMI-8226 | Leukemia | Cancer | WT | Negative | Pass |
| UO-31 | Renal | Cancer | NA | Negative | Pass | RXF_393L | Renal | Cancer | WT | Negative | Pass |
| BT142 | CNS | Cancer | WT | ALT | Pass | SK-MEL-28 | Melanoma | Cancer | WT | Negative | Pass |
| GBM14 | CNS | Cancer | WT | ALT | Pass | SK-MEL-5 | Melanoma | Cancer | WT | Negative | Pass |
| Hs_729T | Muscle | Cancer | WT | ALT | Pass | SN12C | Renal | Cancer | WT | Negative | Pass |
| JW40_frozen | Prostate | Cancer | WT | ALT | Pass | SNU-182 | Liver | Cancer | WT | Negative | Pass |
| U2OS | Bone | Cancer | WT | ALT | Pass | SNU-449 | Liver | Cancer | WT | Negative | Pass |
| Caco-2 | Colon | Cancer | WT | NA | NA | SR | Leukemia | Cancer | WT | Negative | Pass |
| Caki-1 | Renal | Cancer | WT | NA | NA | SW620 | Colon | Cancer | WT | Negative | Pass |
| HeLa | Cervical | Cancer | WT | NA | Pass | T-47D | Breast | Cancer | WT | Negative | Pass |
| Jurkat | Leukemia | Cancer | WT | NA | NA | TK-10 | Renal | Cancer | WT | Negative | Pass |
| LoVo | Colon | Cancer | WT | NA | NA | VCaP | Prostate | Cancer | WT | Negative | Pass |
| MIA_PaCa-2 | Pancreas | Cancer | WT | NA | NA | VCaP-VCR | Prostate | Cancer | WT | Negative | Pass |
| NB4 | Leukemia | Cancer | WT | NA | NA | | | | | | |
| SK-BR-3 | Breast | Cancer | WT | NA | Pass | | | | | | |
| SW480 | Colon | Cancer | WT | NA | Pass | | | | | | |
| TCam-2 | Seminoma | Cancer | WT | NA | Pass | | | | | | |

[a] Cells types include primary, immortalized or cancer. Imm_HPV indicates an HPV immortalized cell line. Imm_SV40 indicates an SV40 immortalized cell line. Imm_TERT indicates that the cell line was immortalized b overexpressing the catalytic subunit of telomerase, *TERT*.

[b] Cell lines with *TERT* promoter mutations are indicated as “Mut” for mutant or “WT” for wild type.

[c] Cell lines that employ the telomerase-independent telomere maintenance mechanism, Alternative lengthening of Telomeres (ALT), are indicated as “ALT”.

[d] Authentication of human cell lines using STR DNA profiling.

### Table S2. Primers used to prepare bisulfite sequencing amplicon libraries

| Amplicon | Strand | Amplicon Size (bp) | Coordinates (Hg19/GRCh37) | Forward Primer (5'-3') | Reverse Primer (5'-3') |
|---|---|---|---|---|---|
| **Tert_1+** | plus | 272 | 1295525 1295796 | GGTTTAGGTTGTGGGGTAATT | AAAAAATATTACAAAAAAACACTCC |
| **Tert 14** | plus | 248 | 1291967 1292214 | GATTAGAGAATTTAAATTTTTTTTAA | CATAAAATTAACACTCCTAACAC |
| **Tert 15** | plus | 162 | 1292313 1292474 | GGGAGTTTGTGGGGAGGGGGTGAA | TCAAAAAATAATACTACTAAACCCTAC |
| **Tert 16** | plus | 144 | 1292603 1292746 | TTATTTTGTTTTGGTATAATTAA | CTAAAACAAAAAAATCACTTAAACCC |
| **Tert 17** | plus | 253 | 1292879 1293131 | TAGTTTTTTAAAGTGTTGGGA | ACAATAAAAAAAAATATCTAAAAACAC |
| **Tert 18** | plus | 254 | 1293101 1293354 | GTTTGTGTTTTTAGATATTTTTTTTTA | CAAAACCCTAATCCTCCTATCTCCAT |
| **Tert 19** | plus | 262 | 1293381 1293642 | GTATTTAGTTTTGGGGTTTGGGT | CCAACACAACAACCCCTAACAAAT |
| **Tert_2-** | plus | 297 | 1293895 1294191 | TTGGAATTTAGAAAGATGGTTTTTA | CTATATAATATCACCTACCAAACCC |
| **Tert 20** | plus | 213 | 1293618 1293830 | TATTTGTTAGGGGTTGTTGTGTTG | CCCCTATTTCTAAAACTACTTAAAAAC |
| **Tert 21** | plus | 214 | 1293808 1294021 | TTAAGTAGTTTTAGAAATAGGGG | AAACCAAACACTTCCTCTACTCCTC |
| **Tert 22** | plus | 202 | 1293996 1294197 | TGAGGAGTAGAGGAAGTGTTTGGTT | TAATTTCTATATAATATCACCTACC |
| **Tert 23** | plus | 197 | 1294173 1294369 | GGTAGGTGATATTATATAGAAA | AAATCCCCCTAAACCTACCAACCC |
| **Tert 25** | plus | 247 | 1294538 1294784 | GTGTTAGTAGGTGAATTAGTA | CAAATATCCTACCTAAAAAAACTAATAAC |
| **Tert 26** | plus | 255 | 1294754 1295008 | GGGTTATTAGTTTTTTTAGGTAGGATA | CCCCAAAACTAACRACTAATACAAC |
| **Tert 27** | plus | 201 | 1294934 1295134 | GTATATTAGGTATTGGGTTATTA | TAAAAAACCCTAACCCCRACCACCC |
| **Tert 28** | plus | 228 | 1294996 1295223 | GTTAGTTTTGGGGTTTTAGG | AACCCTCCCAACCCCTCCCCTTCCTT |
| **Tert 29** | plus | 207 | 1295197 1295403 | AAAGGAAGGGGAGGGGTTGGGAGGG | AAACCAAACCRAACTCCCAATAAAT |
| **Tert_2b** | minus | 218 | 1293882 1294162 | AAGAAGTTATTTTTTTGGAGGGTG | TAACATCCAAAACCTAAAACCCA |
| **Tert 30** | plus | 257 | 1295314 1295570 | GGGTGTTYGGGTTTTTAGTTTTT | CCTCCACATCATAACCCCTCCCTC |
| **Tert 31** | plus | 242 | 1295555 1295796 | GGTTATGATGTGGAGGTTTTGGGAATAGG | AAAAAATATTACAAAAAAACACTCC |
| **Tert 32** | plus | 229 | 1295716 1295944 | GGTTGGGGATGAATTYGAGGA | CCTAACTCCATTTCCCACCCTTTCT |
| **Tert 33** | plus | 231 | 1295919 1296149 | GAGAAAGGGTGGGAAATGGAGTTAG | CTTCTACTACTAAACTAAAAATC |
| **Tert 34** | plus | 209 | 1296127 1296335 | GATTTTTAGTTTAGTAGTAGAAG | TTTATTAACATTTCAATATTTACC |
| **Tert 35** | plus | 241 | 1296283 1296523 | GGTTTTGTAGGGATGTTGTAGTTGAGG | CTTAAAAATCACTAAAAAAATTTCT |
| **Tert 37** | minus | 257 | 1296158 1296414 | GTTTAAATGTTAGTTTTATAAATAAAG | CCACTAATCCCCTCCAAACCT |
| **Tert 38** | minus | 215 | 1296018 1296232 | GGAGTTTGGATTTTTGGGAAGTTTTTAG | CTAACAAATAAAACCAACATCTAATCAC |
| **Tert 39** | minus | 213 | 1295826 1296038 | GATGTTGGTTTTATTTGTTAGATAGAG | TCATTTCTCTTTACAAATTCTCA |
| **Tert 40** | minus | 231 | 1295621 1295851 | GTTTGAGAATTTGTAAAGAGAAATG | AAACCCAAAACTACCTCCAAATCC |
| **Tert 41** | minus | 172 | 1295473 1295644 | GGATTTGGAGGTAGTTTTGGGTTT | CTCCCAAAATACAAAAACRCCAAC |
| **Tert 42** | minus | 209 | 1295378 1295586 | GTATTTGTTTTTAGGGTTTTTATATTATG | AATCCACTAAAAACCCRACCTAACCC |
| **Tert 43** | minus | 205 | 1295292 1295496 | GTTGGYGTTTTTGTATTTTGGGAG | CTAAAAAATAAAAAAACAAAACRAATACC |
| **Tert 44** | minus | 235 | 1295109 1295343 | GTGGYGGAGGGATTGGGGATT | AAAATAACCRAAACCAAAACTTCCC |
| **Tert 45** | minus | 255 | 1294985 1295239 | GGTTTAGTTTTTTTYGGGTTTTTTTAG | CTACACCAACCRCCAACCCTAAAACCCC |
| **Tert 47** | minus | 247 | 1294539 1294785 | GTAGGTGTTTTGTTTGAAGGAGTTGGTGG | TACCAACAAATAAACCAACAC |
| **Tert 48** | minus | 211 | 1294349 1294559 | GTGTTGGTTTATTTGTTGGTA | CTAACAAACCCAAAAAAACCCC |
| **Tert 49** | minus | 232 | 1294139 1294370 | GGGGTTTTTTTGGGTTTGTTAG | CACCCTCCAAAAAAATAACTTCTT |
| **Tert 52** | minus | 216 | 1293616 1293831 | GTTTTTGTTTTTGGAGTTGTTTGGGAA | TACACCTACCAAAAACTACTATACT |
| **Tert 53** | minus | 231 | 1293413 1293643 | GTTAGTATAGTAGTTTTTGGTAGGTGTA | CCACCACCTCCTCACCTAAACTCCT |
| **Tert 55** | minus | 221 | 1293156 1293376 | GGGGTTTAGAAAAGGGGGTAGGTAGAG | CCCAACCTCCTCTATTCACTACTCTAACC |
| **Tough_tert_4** | minus | 183 | 1294348 1294530 | GTTTTTTGTGTTGGTGGTTTTTA | ACTAACAAACCCAAAAAAACCCC |
